# Uncovering High-Order Epistatic Interactions in GWAS via a Machine Learning-Based Feature Engineering Framework

**DOI:** 10.64898/2026.08.03.742638

**Authors:** Jinyoung Byun, Dheeman Saha, Younghun Han, Vikram R. Shaw, Katherine Siminovitch, Christopher I. Amos

## Abstract

**Background:** Genome-wide association studies (GWAS) often fail to identify higher-order epistatic interactions that contribute to complex inheritance patterns of traits and diseases. While machine learning (ML) can capture non-linear relationships, extracting interpretable insights from these models remains a challenge. We propose a novel tree-based feature engineering framework that uses Classification and Regression Trees (CART) to explicitly encode high-order interaction decision paths as dummy variables. We investigate three path-based encoding strategies: (i) all decision paths, (ii) leaf-node paths only, and (iii) internal-node paths only. This approach aims to transform complex decision boundaries into discrete features that capture nonlinear interactions that are not readily captured by traditional association models.

**Results:** The framework was evaluated using genetic data for ANCA-associated vasculitis (AAV). To manage the high dimensionality of the engineered feature space, we applied a comprehensive suite of ML methods across three tasks: (1) Ensemble Learning (Random Forest, XGBoost, and Gradient Boosting Machine); (2) Decision Tree Analysis (CART); and (3) Regression and Classification Tasks (Regularized Linear Regression/LASSO, Support Vector Machine, and Logistic Regression). Stepwise feature selection and regularization were employed to isolate the most informative interaction patterns. Results indicate that incorporating CART-derived interaction paths—particularly those from high-impact regions of the tree—significantly improves classification accuracy and model interpretability compared to using the original feature space alone.

**Conclusions:** The proposed framework provides a robust, scalable methodology for identifying high-order genetic interactions. By bridging the gap between the predictive power of ensemble ML and the necessity for mechanistic insight, this approach offers a clearer mapping of the combinatorial genetic processes underlying complex diseases. While applied here to AAV, the method is highly adaptable for exploring the genetic architecture of diverse populations and complex traits.

## 1 Introduction

Genome-wide association studies (GWAS) have become an important approach for identifying genetic variants associated with complex diseases. By evaluating large numbers of single nucleotide polymorphisms (SNPs), GWAS can reveal genomic regions associated with disease susceptibility and provide insight into biological mechanisms. However, many complex diseases are not driven by single genetic variants alone. Instead, disease risk may reflect nonlinear relationships, correlated markers, and higher-order interactions among multiple variants. These patterns are difficult to detect using conventional single-marker association analysis, especially when genomic datasets contain far more features than samples.

This challenge is particularly important in rare and heterogeneous diseases, where sample sizes are often limited, and disease mechanisms may differ across clinical sub-types. ANCA-associated vasculitis (AAV) is one such disease context. AAV includes clinically distinct subtypes, such as granulomatosis with polyangiitis (GPA) and microscopic polyangiitis (MPA), and prior genetic studies have identified susceptibility loci associated with disease risk. However, identifying multivariate genomic patterns that distinguish disease groups or subtypes remains difficult because GWAS data are high-dimensional, sparse, and biologically complex.

A major unresolved problem in GWAS-based prediction is determining how to model complex SNP relationships while maintaining interpretability. Many computational models can be applied to genomic classification, but their performance can vary depending on the algorithm, feature representation, optimization strategy, and outcome definition. In addition, models that improve prediction may not necessarily provide interpretable genetic patterns, while simpler models may fail to capture non-linear interactions. This creates a trade-off between predictive accuracy, computational efficiency, and biological interpretability.

Another challenge is the lack of systematic comparison across machine learning strategies in the same GWAS setting. Without a consistent evaluation framework, it is difficult to determine whether observed performance differences are due to the classifier itself, the optimization procedure, or the way genomic features are represented. This is especially relevant when the goal is not only to classify disease status but also to identify informative SNP patterns that may reflect underlying genetic interactions.

Therefore, there is a need for a comparative framework that evaluates machine learning models for GWAS-based disease classification while also examining interpretability and computational cost. Such a framework is important for understanding which modeling strategies are most suitable for high-dimensional biomedical data and for determining whether interaction-oriented feature representations can support the discovery of disease-relevant genomic patterns. In this study, we address this problem using GWAS data from AAV as a representative complex disease setting.

## 2 Background

The increasing availability of high-dimensional biomedical data has catalyzed the use of machine learning (ML) techniques in complex disease research [5]. Compared to traditional statistical approaches, ML methods offer greater flexibility in handling non-linearity, interactions, and noisy or correlated variables—common characteristics of genomic datasets such as those generated by genome-wide association studies (GWAS) [9]. However, the diversity of ML algorithms, each with unique strengths and inductive biases, poses a challenge in selecting the most appropriate method for a given task [9]. As a result, systematic comparisons of ML approaches are essential for guiding optimal method selection and improving predictive modeling in biomedical data mining. Among the most widely used ML methods are ensemble learning techniques, including Random Forest (RF) [2], Gradient Boosting Machines (GBM) [8], and eXtreme Gradient Boosting (XGBoost) [4], which improve predictive performance by aggregating the outputs of multiple decision trees to mitigate overfitting. In contrast, single decision-tree methods like Classification and Regression Trees (CART) [3] offer interpretable, rule-based decision structures, while regularized regression techniques such as the Least Absolute Shrinkage and Selection Operator (LASSO) [10], Support Vector Machine (SVM) [1], and Logistic Regression models [6] offer parsimonious solutions that are particularly useful when the number of features exceeds the number of observations.

Evaluating these diverse ML approaches in a comparative framework provides insight into their relative performance across different modeling objectives, such as classification accuracy and feature selection, especially in complex genetic data. While several susceptibility loci have been identified through traditional GWAS [7], these methods often overlook intricate gene-gene interactions and non-additive effects. To bridge this gap, we propose a novel ML-based feature engineering framework that explicitly targets high-order interaction decision paths learned by a CART classifier. By extracting these paths and encoding them as dummy variables—derived from all nodes (i.e., considering all paths), leaf nodes, or internal nodes—we transform complex, nonlinear feature interactions into a structured format compatible with various classification models. To demonstrate the utility of this approach in a comprehensive genetic research context, we applied the framework to GWAS data from patients with ANCA-associated vasculitis (AAV) [7]. AAV comprises a group of rare autoimmune diseases, including granulomatosis with polyangiitis (GPA) and microscopic polyangiitis (MPA), characterized by small-vessel inflammation and the presence of antineutrophil cytoplasmic antibodies (ANCA) [7]. By applying a comparative ML framework to AAV, we aim to assess the relative performance of diverse models that integrate path-based feature engineering and to uncover novel genetic interactions associated with disease-relevant genomic patterns.

## 3 Methods

This study employed a comparative machine-learning framework to evaluate classification performance and feature interpretability using genomic and clinical data from the study population. The methodological workflow included data preprocessing, model optimization, feature selection, and systematic comparison across multiple learning architectures. We evaluated ensemble and boosting models, linear and regularized classifiers, and a single CART-based decision-tree architecture to capture both complex nonlinear patterns and interpretable decision boundaries. In addition, a CART-driven feature engineering pipeline was used to transform decision paths into derived features, enabling downstream classifiers to assess whether tree-based rule structures improved predictive performance and supported the identification of meaningful feature interactions.

### 3.1 Data Source and Study Population

The dataset used in this study was obtained from GWAS of ANCA-associated vasculitis (AAV) previously described by Merkel et al.[7]. The cohort comprises patients of European ancestry diagnosed with Granulomatosis with Polyangiitis (GPA) and Microscopic Polyangiitis (MPA), as well as healthy controls. To extend the original study’s focus on independent risk loci, we used quality-controlled genotype data to examine higher-order epistatic interactions. Single-nucleotide polymorphisms (SNPs) were encoded using both additive and dominant genetic models, a strategy designed to capture a broad spectrum of inheritance patterns, including heterozygote effects that may be obscured in standard univariate analyses.

### 3.2 Model Optimization and Feature Selection Strategies

To ensure a robust comparison across diverse algorithmic architectures, we evaluated six distinct classifiers through four integrated optimization paradigms. In all scenarios, model performance was validated using stratified cross-validation, a necessary precaution to preserve the integrity of the results given the AAV dataset’s inherent class distribution. We initially employed Brute-Force Enumeration to conduct an exhaustive evaluation of predefined parameter configurations, ensuring total reproducibility and methodological fairness. To further refine these models, we implemented Grid Search (Cross-Validation) using GridSearchCV to identify the optimal balance between model bias and variance within each classifier’s parameter space. Recognizing the challenges of high-dimensional genomic data, we integrated Forward Selection into our workflow. This greedy iterative procedure begins with an empty feature set and sequentially adds variables that yield the greatest marginal gain with respect to a chosen performance metric, such as the Area Under the Receiver Operating Characteristic curve (AUC) or the F1-score. Finally, we introduce our proposed framework, CART-Driven Interaction Encoding. This approach targets classifiers that may underperform in linear feature spaces by decomposing a trained CART into its constituent decision paths. By transforming hierarchical logic into a high-dimensional dummy variable space, we bridge the gap between raw genomic data and complex disease phenotypes.

### 3.3 Ensemble and Boosting Architectures

Our analysis leveraged the predictive power of ensemble methods, specifically Random Forest (RF) (Supplementary Table A5), XGBoost (Supplementary Table A6), and Gradient Boost (GBM) (Supplementary Table A7). For the Random Forest classifier, we explored a configuration space of 100-500 trees, constraining tree depth to 4 or 6 to mitigate overfitting, and tested various splitting criteria to manage tree complexity. In boosting frameworks, we balanced training speed with stability by varying the learning rate between 0.05 and 0.1 and maintaining moderate tree depths of 3 to 5. To mitigate the risk of memorizing noise in sparse genomic predictors, we employed stochastic subsampling of both samples and features, thereby maintaining generalizability across diverse genetic markers.

### 3.4 Linear and Regularized Classification Tasks

To provide a parsimonious comparison with ensemble methods, we implemented Least Absolute Shrinkage and Selection Operator (LASSO) (Supplementary Table A1), Logistic Regression (Supplementary Table A2), and SVM (Supplementary Table A3). The LASSO model utilized the SAGA solver to handle the non-smooth L1 penalty, with the regularization strength C spanning a wide continuum from 0.01 to 10.0 to encourage sparsity. For the SVM, we explored linear, RBF (radial basis function), and polynomial kernels to capture various decision boundary geometries. Across these tasks, we addressed potential class imbalance by employing a balanced class-weighting strategy that assigns higher weights to underrepresented AAV subtypes. This ensures that the minority classes receive proportional emphasis during training, preventing the models from biasing predictions toward the majority control groups.

### 3.5 Single Decision-Tree Architecture

To provide an interpretable baseline for comparison with ensemble and regularized machine-learning models, we implemented a single CART-based decision-tree classifier (Supplementary Table A4). Unlike ensemble approaches that aggregate predictions across multiple trees, the single-tree architecture produces a transparent sequence of decision rules that can be traced from the root node to terminal leaf nodes. At each internal node, the model partitions the feature space by selecting the split that best separates the outcome classes, while each terminal node represents a final classification region. This structure allows the learned decision paths to be examined directly and mapped back to the original genomic features, making the model useful for identifying candidate feature interactions and rule-based patterns. To reduce overfitting and maintain generalizability, the tree construction was constrained to balance model complexity with predictive performance. Overall, the single decision-tree architecture served as an interpretable reference model for evaluating whether more complex learning strategies provided additional predictive or explanatory value.

### 3.6 CART-Driven Feature Engineering Pipeline

The final phase of our methodology involved implementing a CART-based framework in Figure 1 to improve the predictive accuracy of the primary classifiers. The procedure began by loading the optimized parameter sets for each training and validation partition. We then used the CART model to generate a hierarchical tree, in which each node and its corresponding decision path were defined as discrete binary dummy variables. This transformation enabled the systematic recording of the specific genetic features used as decision criteria for each split in the tree, as shown in Figure 2. Figure 2 presents the outcome of forward feature selection using dummy variables. The first plot shows that the cross-validation score increases sharply after selecting the first few dummy variables and then gradually stabilizes, indicating diminishing returns from adding more variables. The second plot shows the incremental score improvement at each selection step, where the largest gain occurs at the first step, and later additions contribute only marginal improvements. Overall, these results suggest that a small subset of selected dummy variables captures most of the predictive information, while additional variables provide limited performance gain.

**Fig. 1:**
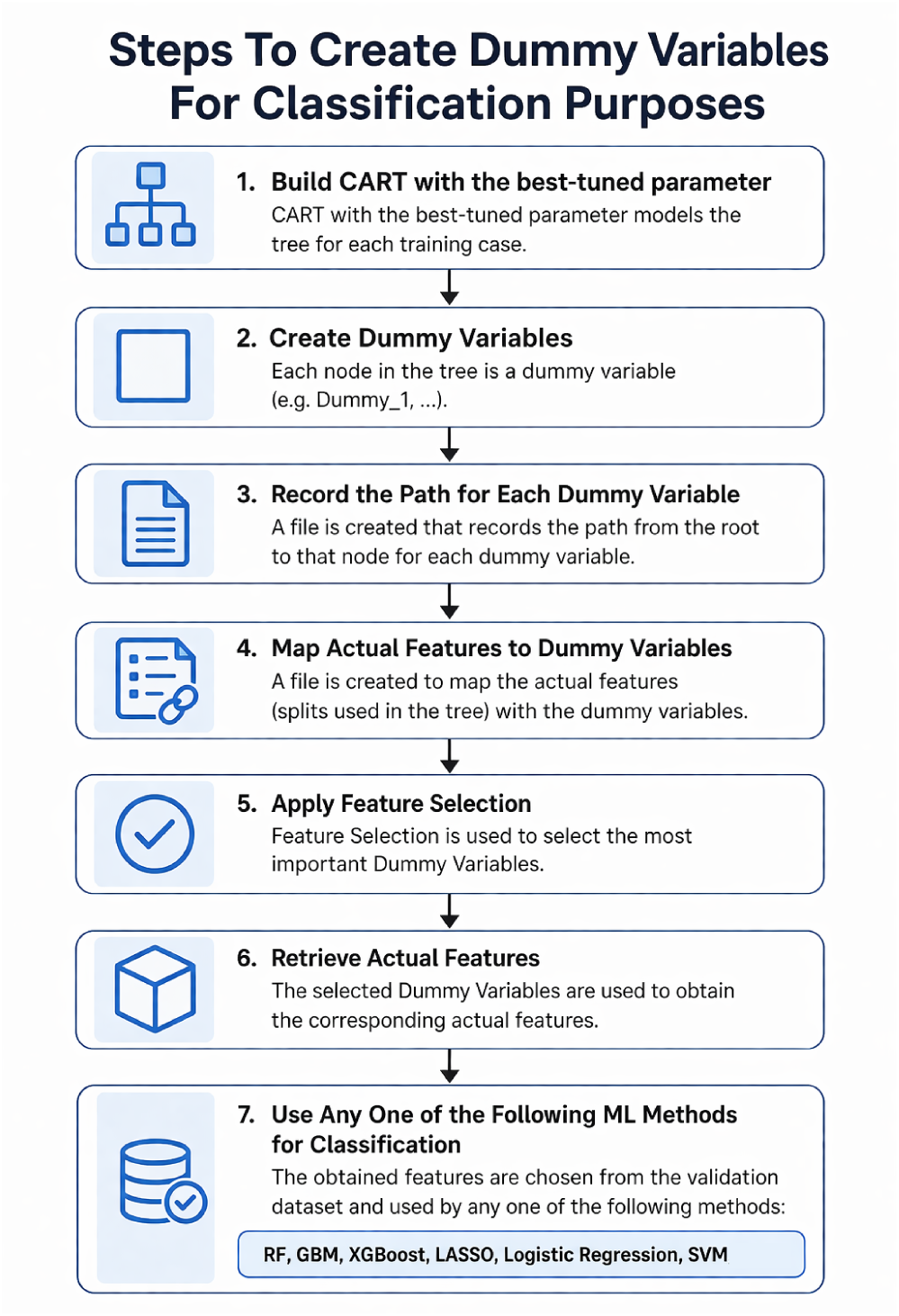
Overview of the Classification Approach using Dummy Variables.

**Fig. 2:**
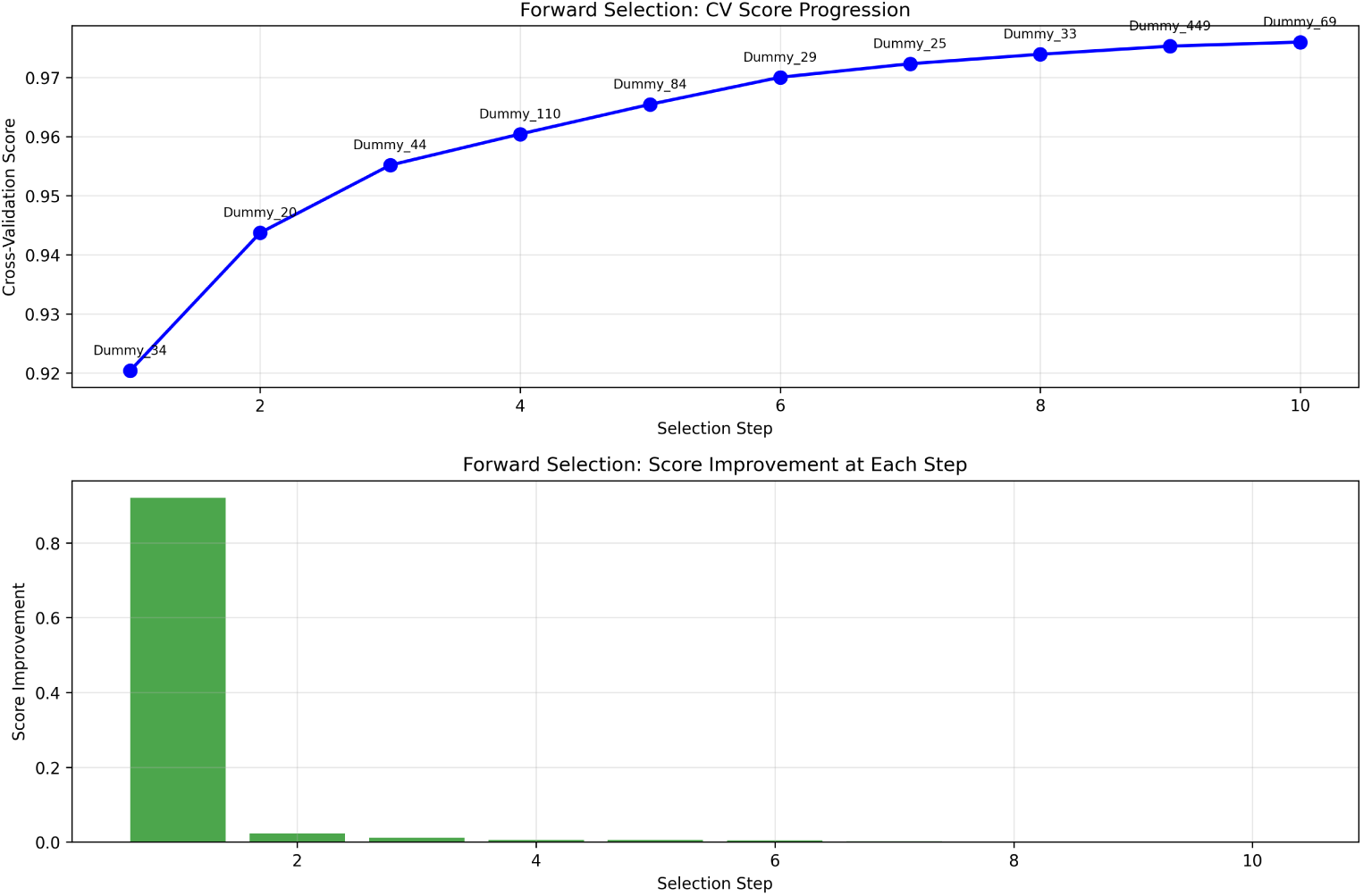
Outcome of stepwise feature selection of the dummy variables for training set 20. The top plot shows the cross-validation scores of the dummy variables. The bottom plot shows the incremental score improvement at each selection step.

Initially, based on Figure 3, the tree is generated, where each node is labeled with the corresponding dummy variable condition. The tree starts from the root node, which contains the full training sample, and applies the first split based on an HLA-related dummy variable. Each subsequent internal node further partitions the data into smaller subgroups using additional dummy-variable rules.

**Fig. 3:**
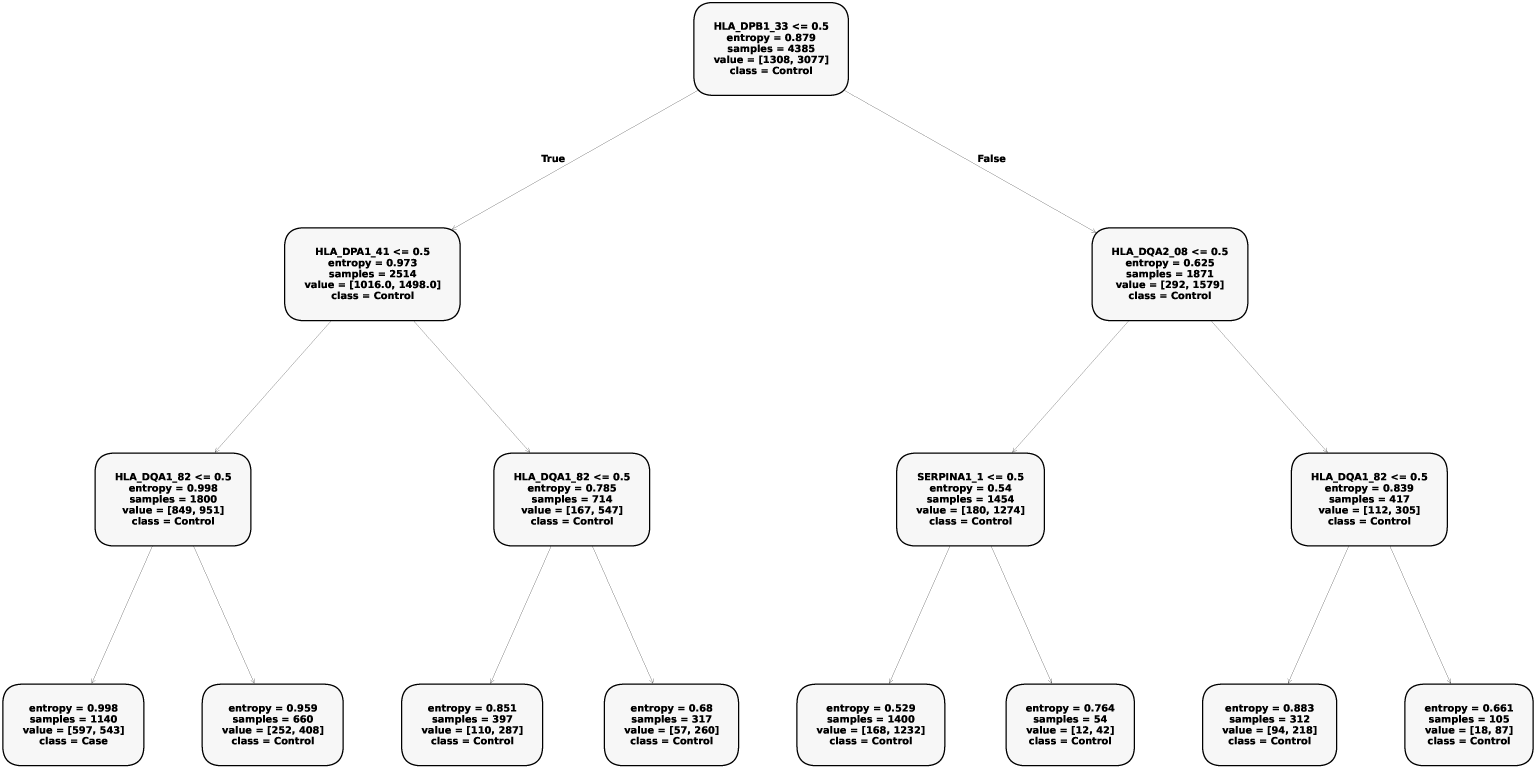
The generated tree after using the CART Classifier based on training set 85. All the nodes are labelled with dummy variables.

The node labels report the splitting condition, entropy, number of samples, class distribution, and predicted class. In this case, most terminal nodes are classified as Control, while one leftmost terminal node is classified as Case, indicating that only a specific decision path identifies a subgroup with stronger case representation. This suggests that the CART model mainly separates the population into control-dominant regions, with a smaller subset of nodes capturing case-associated patterns.

Overall, the figure shows how the CART-derived dummy variables create interpretable decision paths from genetic features to final class labels. The tree also demonstrates that the selected dummy-variable splits provide a structured way to identify important subgroups, but the class distributions remain largely control-dominant across most branches. To evaluate the informational value of different regions within the tree architecture, we investigated three distinct path-extraction strategies as represented in Figure 4. In the first scenario, representing our primary proposed approach, all possible paths from the root node were considered. This comprehensive analysis provided a global view of the interaction space, successfully capturing both broad parental splits and granular terminal interactions. To compare the efficacy of this global model with more localized strategies, we further investigated a subset comprising only paths associated with the leaf nodes, as indicated by the red nodes in Figure 4.

By omitting intermediate segments, this second approach focused exclusively on the classifier’s final decision boundaries.

In contrast, we examined a third strategy that considered only paths that did not terminate at leaf nodes (i.e., internal nodes) as indicated by the blue-colored nodes in Figure 4. By isolating these internal node paths, we were able to evaluate the predictive weight of higher-level, broader genetic splits while intentionally excluding the most granular terminal logic. Each set of generated dummy variables was subsequently subjected to forward stepwise selection, a greedy iterative procedure used to isolate the most predictive subset of features and manage the high dimensionality of the interaction space.

**Fig. 4:**
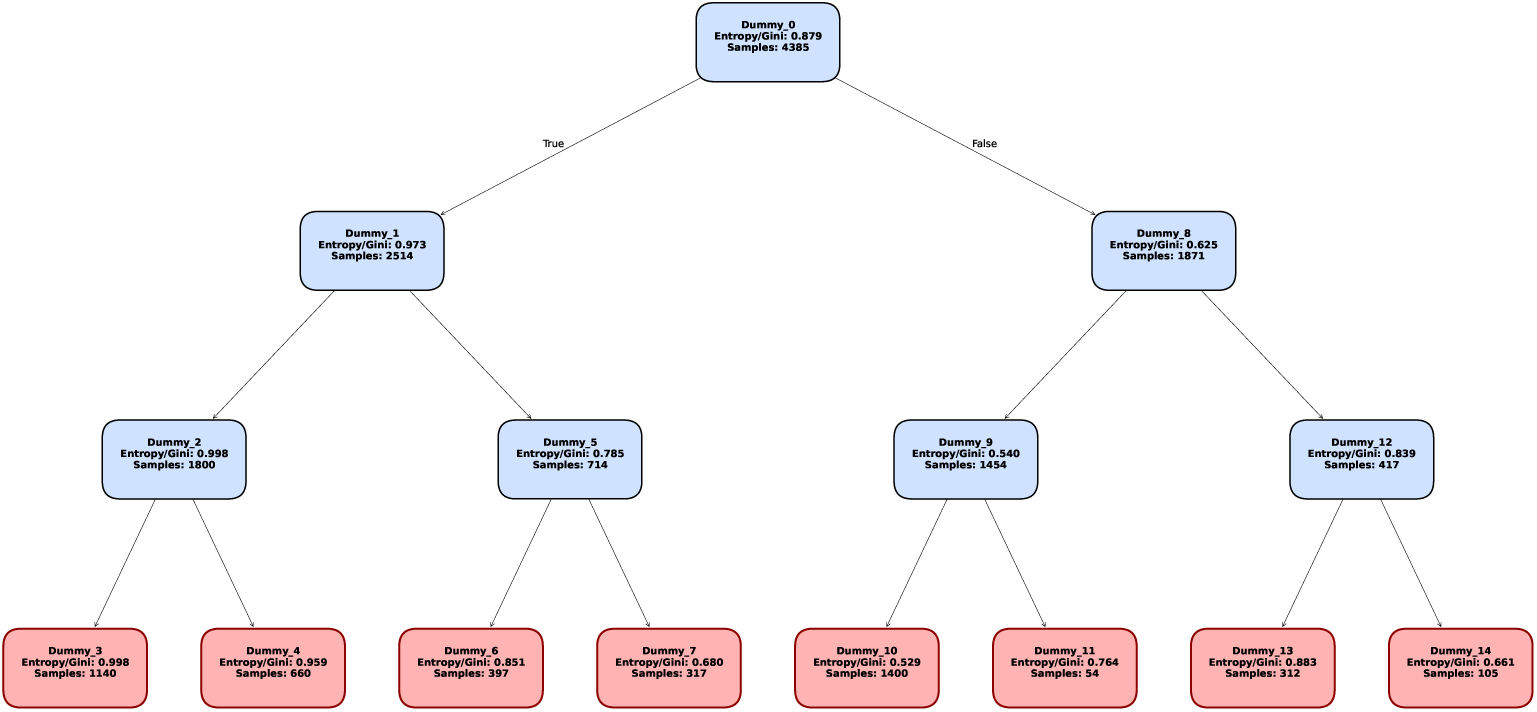
The generated tree using the CART Classifier based on training set 85. The internal nodes are labelled with blue nodes, and the leaf nodes are labelled with red nodes.

Once the optimal dummy variables were identified, they were mapped back to their original genetic features, and their performance was validated using the validation dataset. This systematic comparison allowed us to determine which specific segments of the decision tree—comprehensive, terminal, or internal—captured the most informative high-order genetic interactions underlying AAV susceptibility.

## 4 Results

### 4.1 Comparative Analysis of Classification Accuracy

The performance of each classifier was evaluated by examining the distribution of classification accuracies across multiple iterations, providing insight into the stability, robustness, and predictive strength of each model in Figure 5). Across the ensemble and boosting architectures—comprising RF, GBM, and XGBoost—we observed a consistent trend toward higher stability and accuracy as optimization moved from unguided exploration to feature refinement in Figure 5.a-c. The Brute-Force approach typically yielded broad accuracy distributions, reflecting the models’ sensitivity to hyperparameter variation when configurations are not systematically tuned. In contrast, Grid Search effectively narrowed these distributions, yielding more concentrated results across mid- to high-accuracy ranges by identifying stable learning rates and tree depths. Ultimately, Feature Selection proved the most effective for these ensembles; by isolating a refined subset of informative predictors, the models achieved their highest accuracy peaks with the lowest variance, demonstrating that reducing genomic noise is critical for the generalization of boosted tree models.

**Fig. 5:**
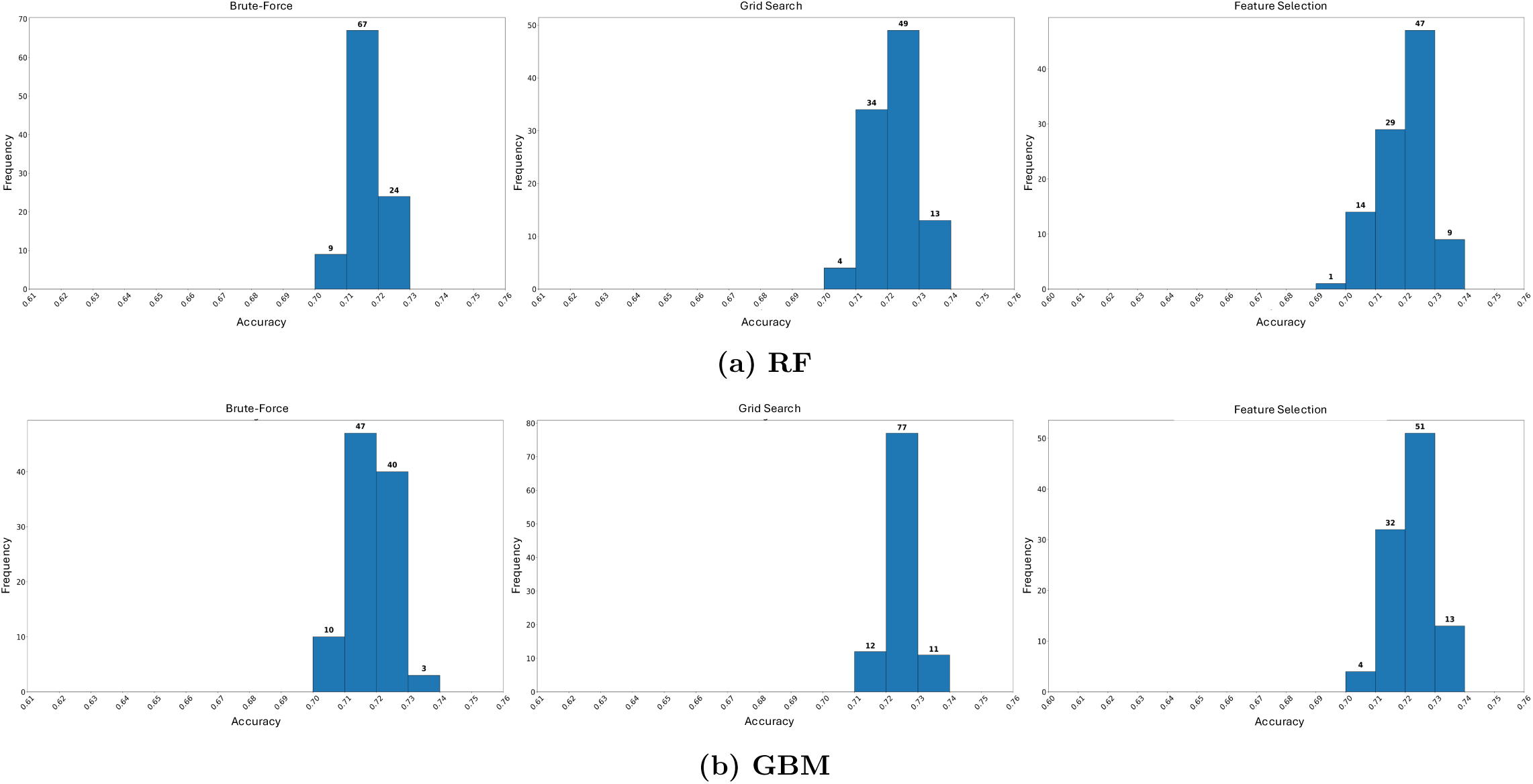
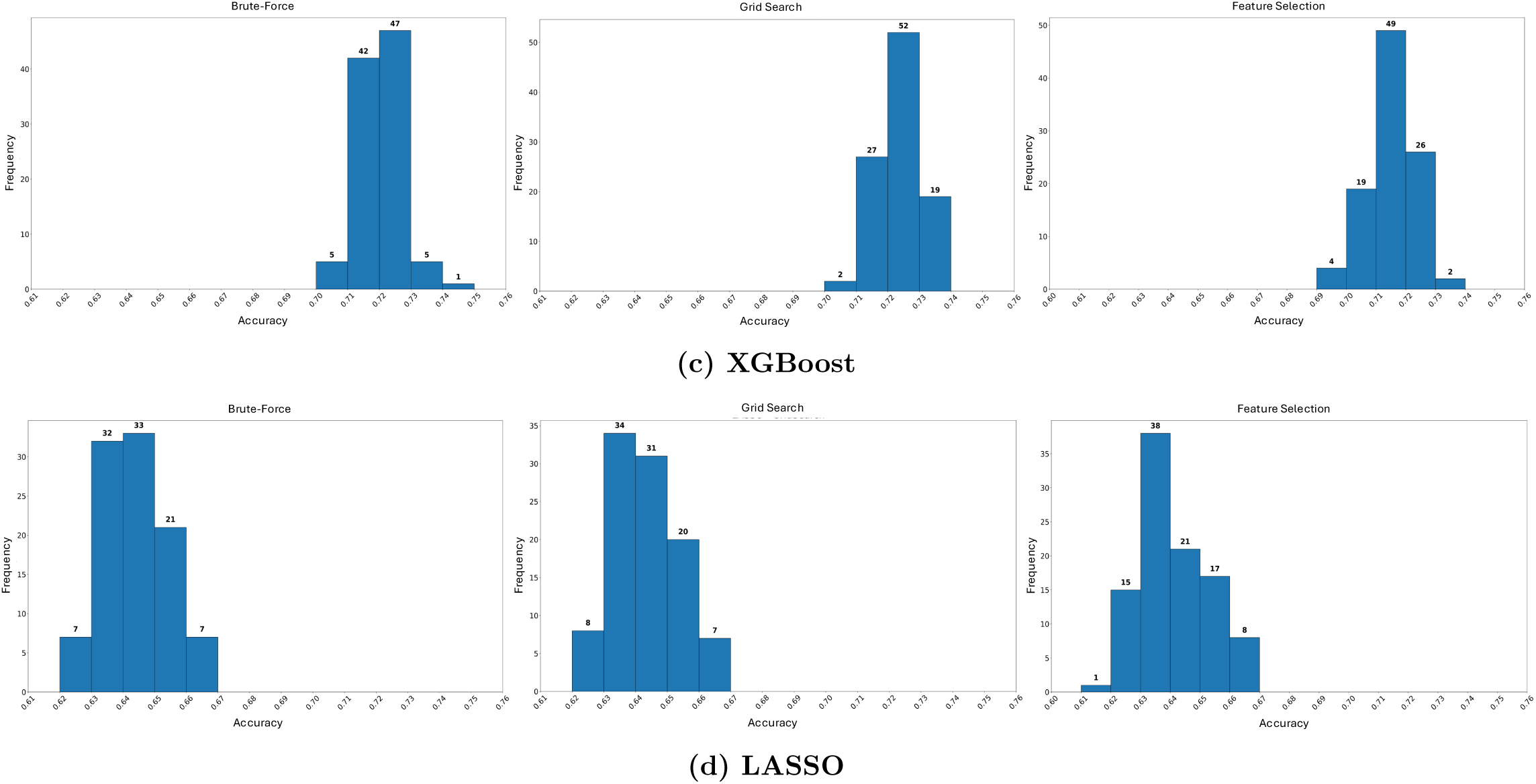
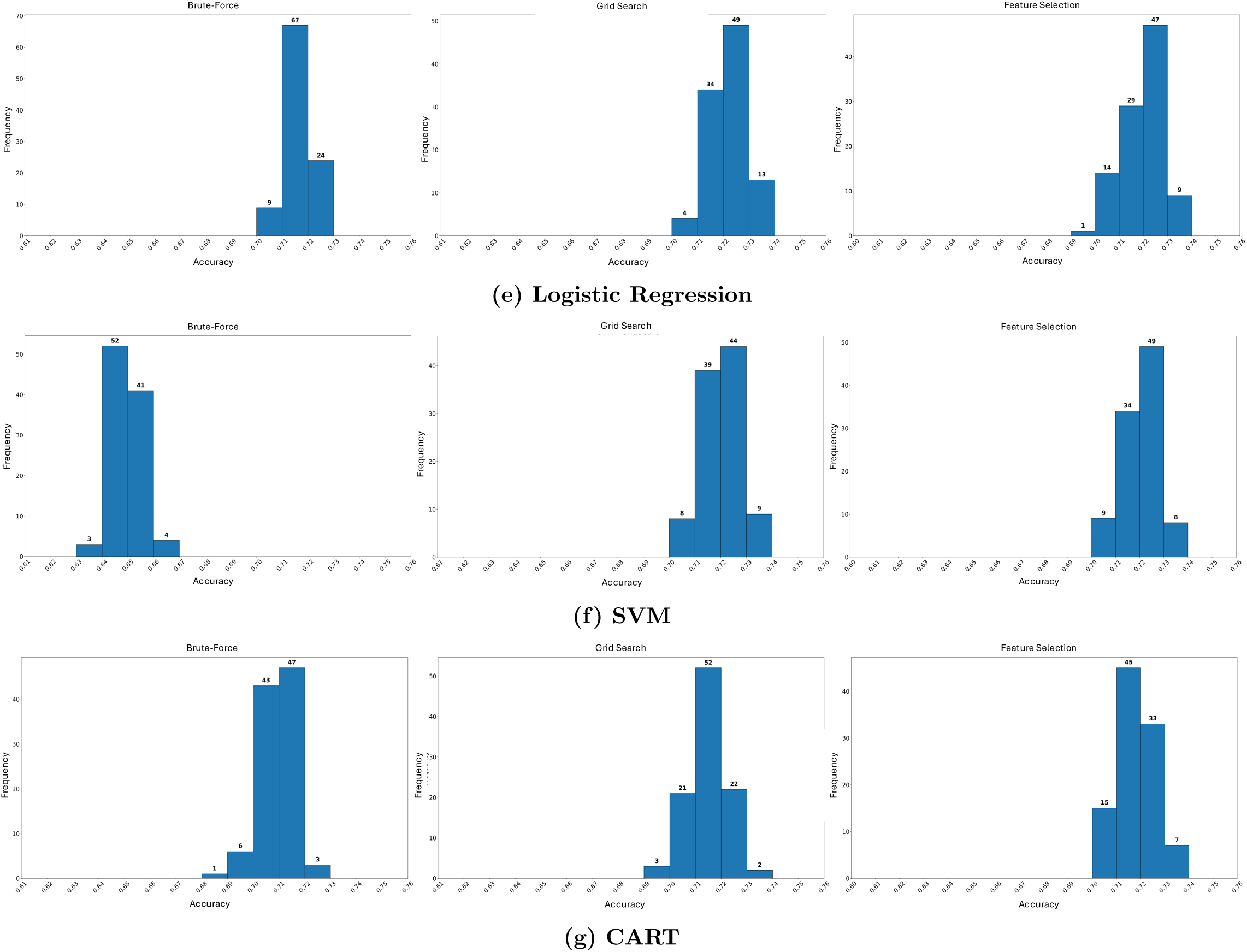
Comparative analysis of classification accuracy using the Brute-Force, Grid Search, and Feature Selection approaches. Panels show: (a) RF, (b) GBM, (c) XGBoost, (d) LASSO, (e) Logistic Regression, (f) SVM, and (g) CART.

The linear and margin-based models, including LASSO, Logistic Regression, and SVM, exhibited a similar evolutionary performance profile in Figure 5.d-f. For LASSO and Logistic Regression, the Brute Force approach often yielded moderate spread, with clusters near the mid-range, because many parameter choices did not meaningfully improve the linear fit. Grid Search provided a significant improvement in consistency, particularly for identifying optimal regularization strengths (C values). For the SVM, which is notably sensitive to high-dimensional noise, the transition to Feature Selection was particularly impactful. By refining the feature set, we observed a distinct shift toward higher accuracy and a more stable decision margin, as the models were better equipped to map the complex genetic boundaries of AAV without interference from redundant variables.

Finally, the CART classifier’s performance highlighted the inherent volatility of single-tree models in Figure 5.g. The Brute Force distribution was the most dispersed among all tested classifiers, confirming CART’s high sensitivity to depth and split thresholds. However, when paired with Feature Selection, CART exhibited a clear concentration of runs reaching high performance. This suggests that, although individual trees are prone to overfitting on noisy genomic data, they become highly effective and interpretable classifiers when the input dimensionality is reduced to the most informative genetic markers.

Through this systematic comparison, it is evident that while hyperparameter tuning via Grid Search improves reliability, the integration of Feature Selection consistently yields the most robust and accurate classification results across all machine learning frameworks investigated in this secondary analysis of AAV GWAS data.

### 4.2 Synthesis of Model Performance and Optimization Impact

The comparative analysis reveals a clear hierarchy in classification efficacy, primarily driven by the underlying model architecture rather than the specific optimization paradigm in Table 1. A notable finding is the superior performance of tree-based ensemble methods—specifically RF, GBM, and XGBoost—which consistently achieved the highest accuracies, ranging from 71.45% to 72.83%. The inherent robustness of these models is evidenced by their stability; for instance, RF maintained a narrow performance band (71.99%-72.33%), indicating that its ensemble nature effectively buffers against hyperparameter variance and minor changes in feature composition.

**Table 1:** Overview of Classification Accuracy using Brute-Force Approach.

| Algorithm | Mean (%) | Medium (%) | Maximum (%) | Minimum (%) |
| --- | --- | --- | --- | --- |
| GBoost | 71.85 | 71.87 | 73.79 | 70.23 |
| LASSO | 64.41 | 64.34 | 66.47 | 62.11 |
| Logistic Regression | 64.44 | 64.35 | 66.62 | 62.09 |
| Random Forest | 71.61 | 71.62 | 72.83 | 70.55 |
| XGBoost | 72.12 | 72.09 | 74.06 | 70.45 |
| SVM | 64.97 | 64.94 | 66.14 | 63.74 |
| CART | 70.90 | 71.00 | 72.72 | 68.92 |

In contrast, linear models such as LASSO and Logistic Regression occupied a distinct mid-range tier, with accuracies spanning 64.01% to 64.75%. The minimal deviation observed across the three experimental settings suggests that these models reached a performance ceiling inherent to their linear assumptions. Interestingly, although Feature Selection often serves as a stabilizer, it resulted in marginal reductions in accuracy for LASSO. This suggests that in the high-dimensional AAV landscape, even features with low individual effect sizes may contribute to a collective signal that regularized linear models rely upon for prediction.

The SVM and CART models demonstrated intermediate but competitive results. SVM performance remained stable near the 72% mark, benefiting slightly from feature refinement, which reflects its sensitivity to high-dimensional noise. CART, however, exhibited a unique sensitivity to optimization; while Brute Force yielded its peak performance (72.33%), the constraints imposed during Grid Search and Feature Selection resulted in slight drops in accuracy. This implies that while pruning and depth constraints improve a tree’s theoretical generalizability, they may also limit the model’s ability to capture the highly granular, complex interactions present in the raw genetic data. Overall, the data suggest that Brute Force and Grid Search yield comparable results across most architectures, whereas Feature Selection serves as a specialized tool that either refines the signal for noise-sensitive models such as SVMs or acts as a regularizer for tree-based models. These patterns underscore that, for AAV GWAS data, the selection of a nonlinear ensemble architecture is the most significant factor in achieving high classification accuracy, outperforming the marginal gains from varying the hyperparameter optimization strategy.

### 4.3 Area Under the Curve Distribution and Discriminative Power

To quantify the classifiers’ ability to distinguish between positive and negative ANCA-associated vasculitis (AAV) instances, we evaluated the Area Under the Receiver Operating Characteristic Curve (AUC). As a threshold-independent metric, AUC provides a robust measure of model performance; values approaching 1.0 indicate superior discriminative ability.

#### 4.3.1 Stability and Variability Across Optimization Settings

The stability of predictive performance was assessed across 100 repeated experiments, revealing distinct patterns in how feature construction and optimization influence model robustness.

For grid search optimization, across all classifiers, systematic hyperparameter tuning consistently yielded the most stable and unimodal AUC distributions. For ensemble methods such as RF in Figure 6(a), GBM (Figure 6(b)), and XGBoost (Figure 6(c)), this approach yielded concentrated spreads with the highest mean AUC values (0.73 for XGBoost). The limited dispersion suggests that systematic tuning of learning rates, tree depths, and estimators effectively buffers the models against variability in random data partitioning.

**Fig. 6:**
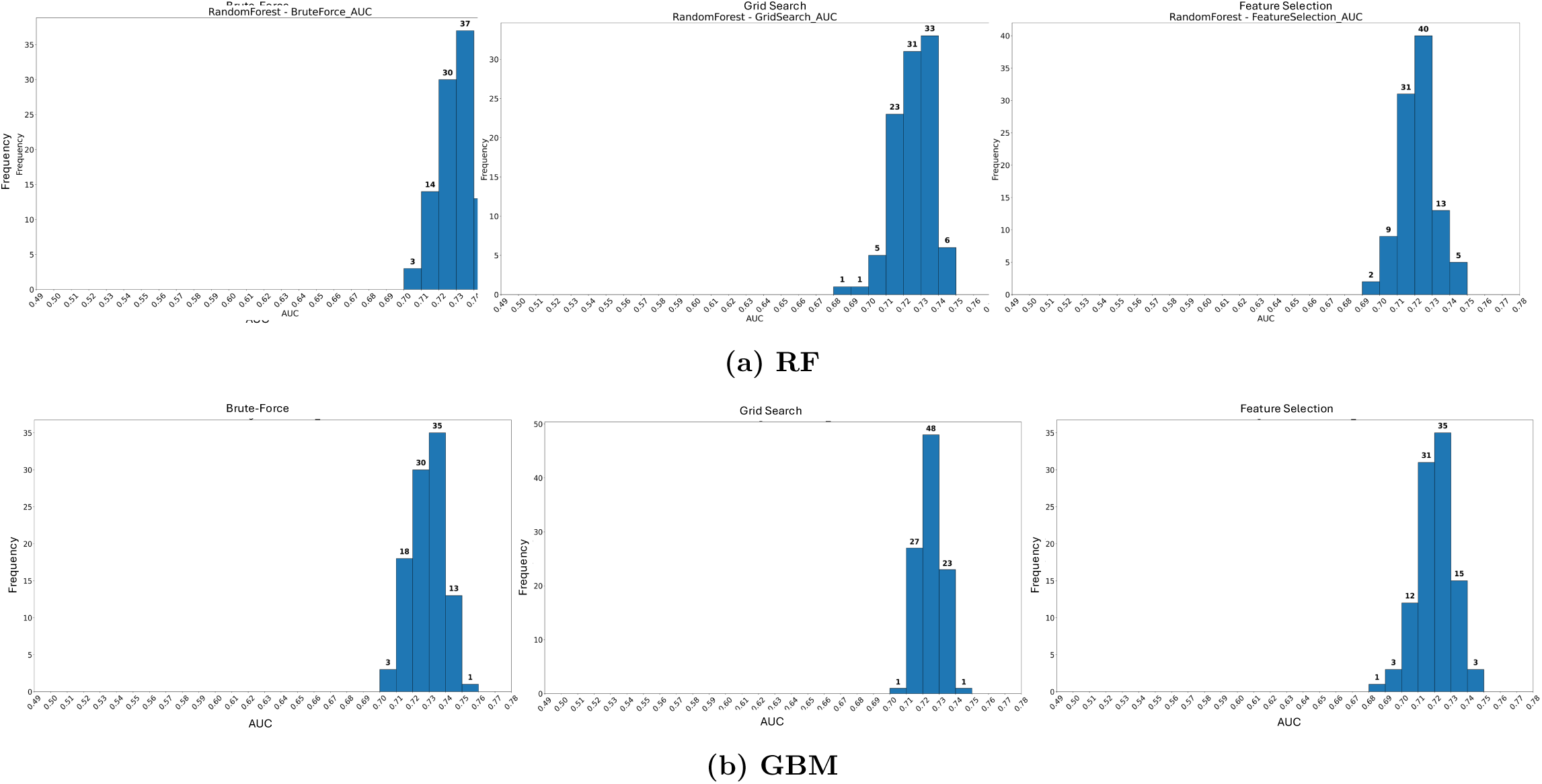
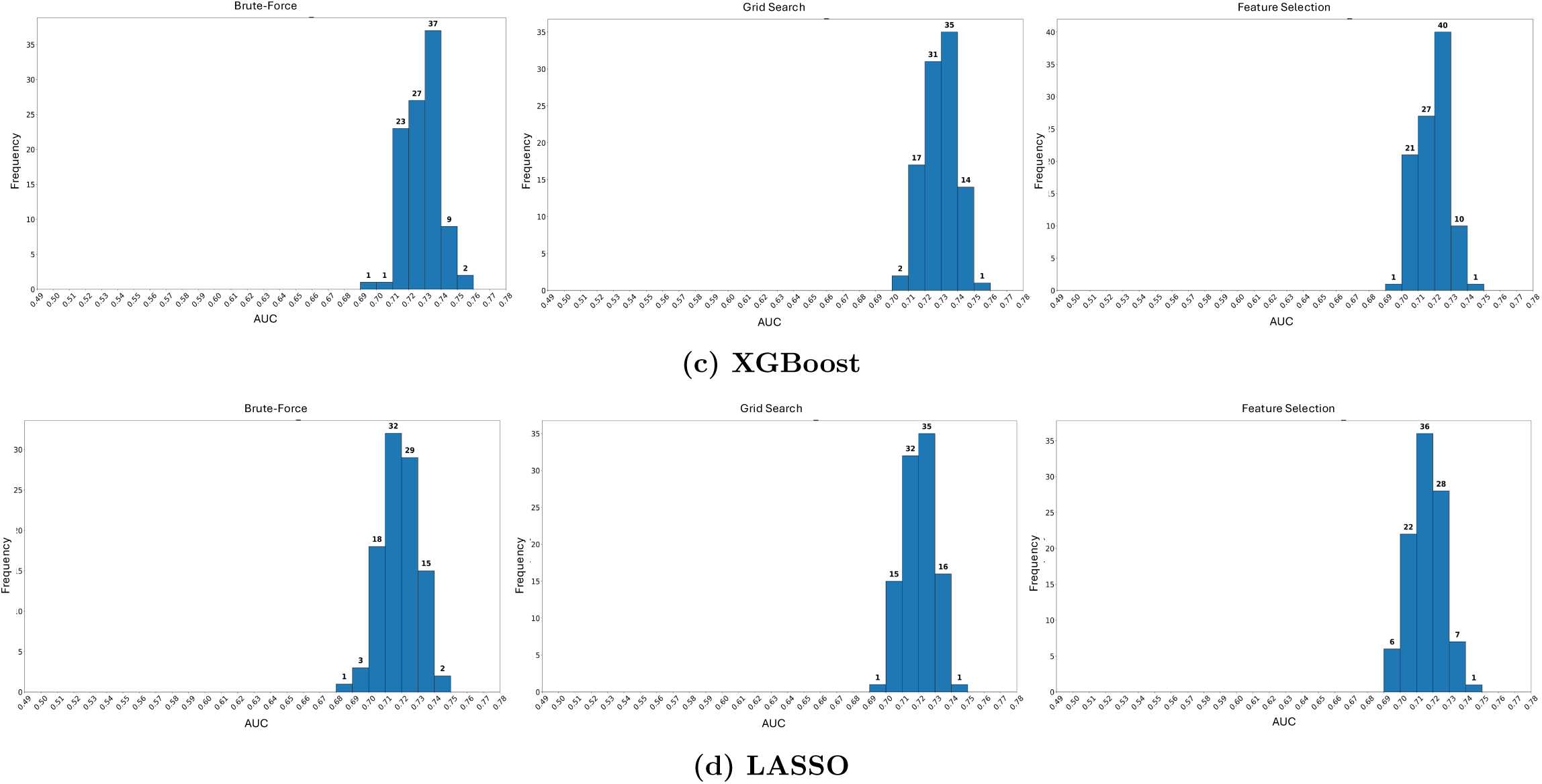
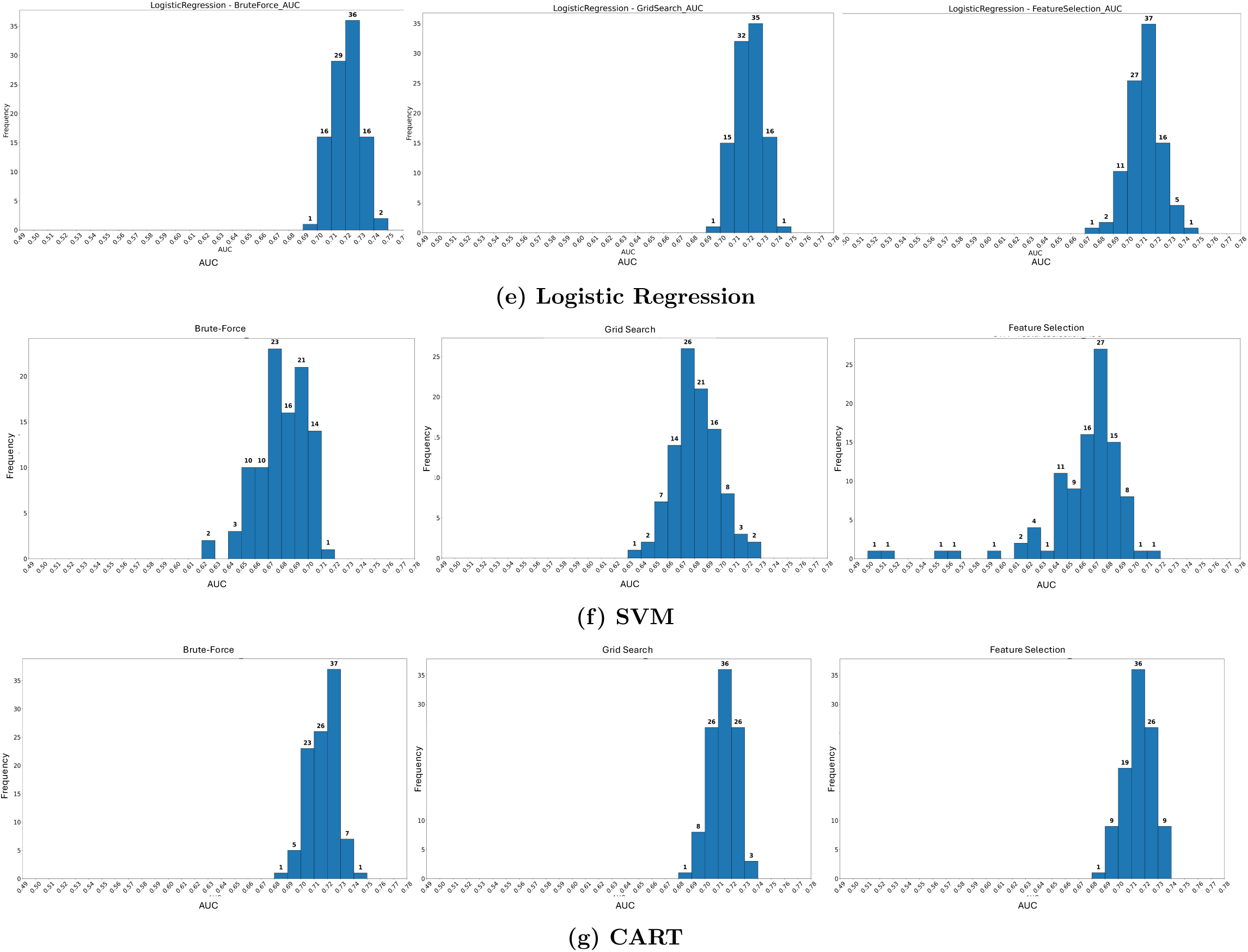
Comparative analysis of AUC using the Brute-Force, Grid Search, and Feature Selection approaches. Panels show: (a) RF, (b) GBM, (c) XGBoost, (d) LASSO, (e) Logistic Regression, (f) SVM, and (g) CART.

**Fig. 7:**
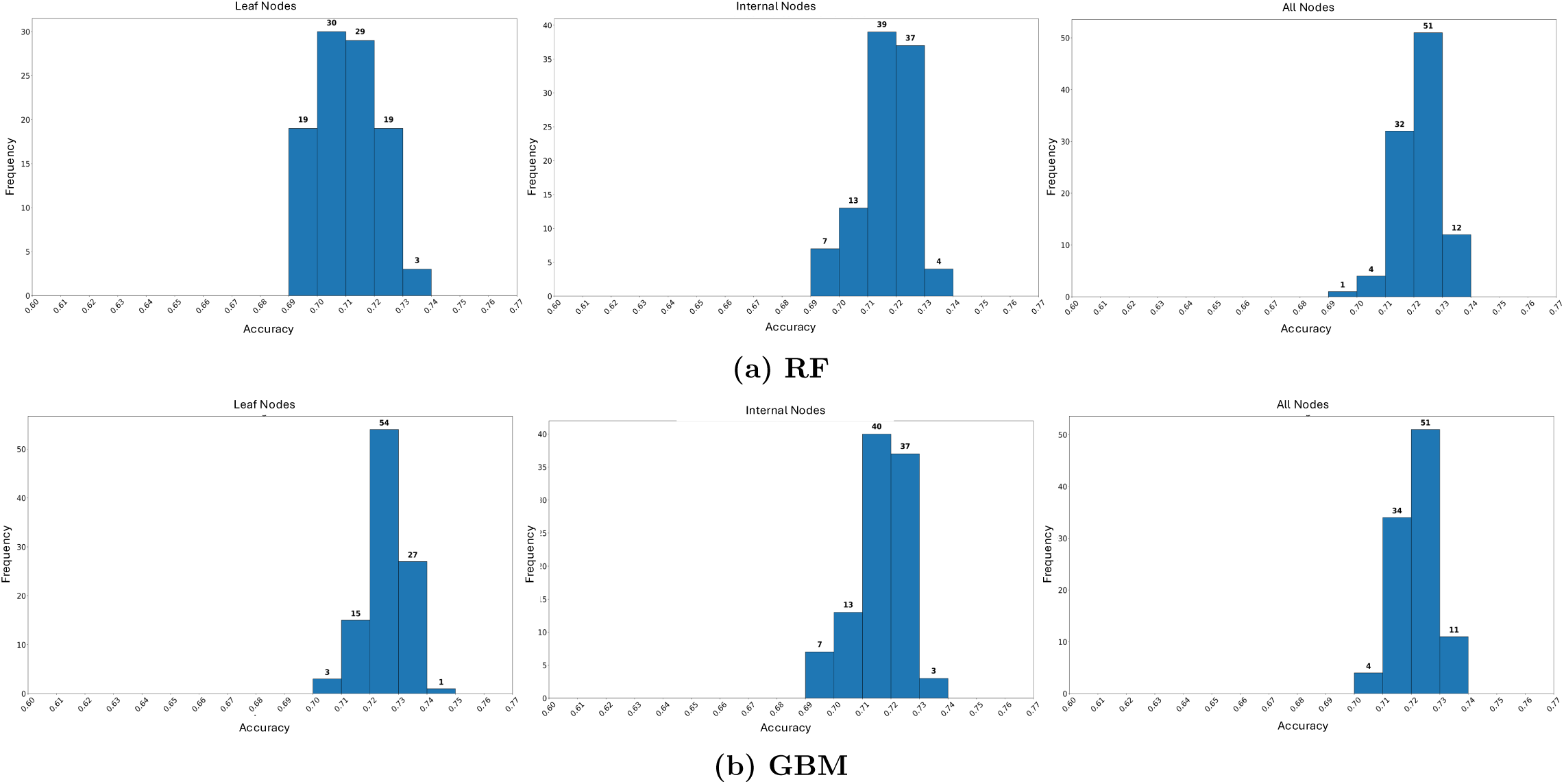
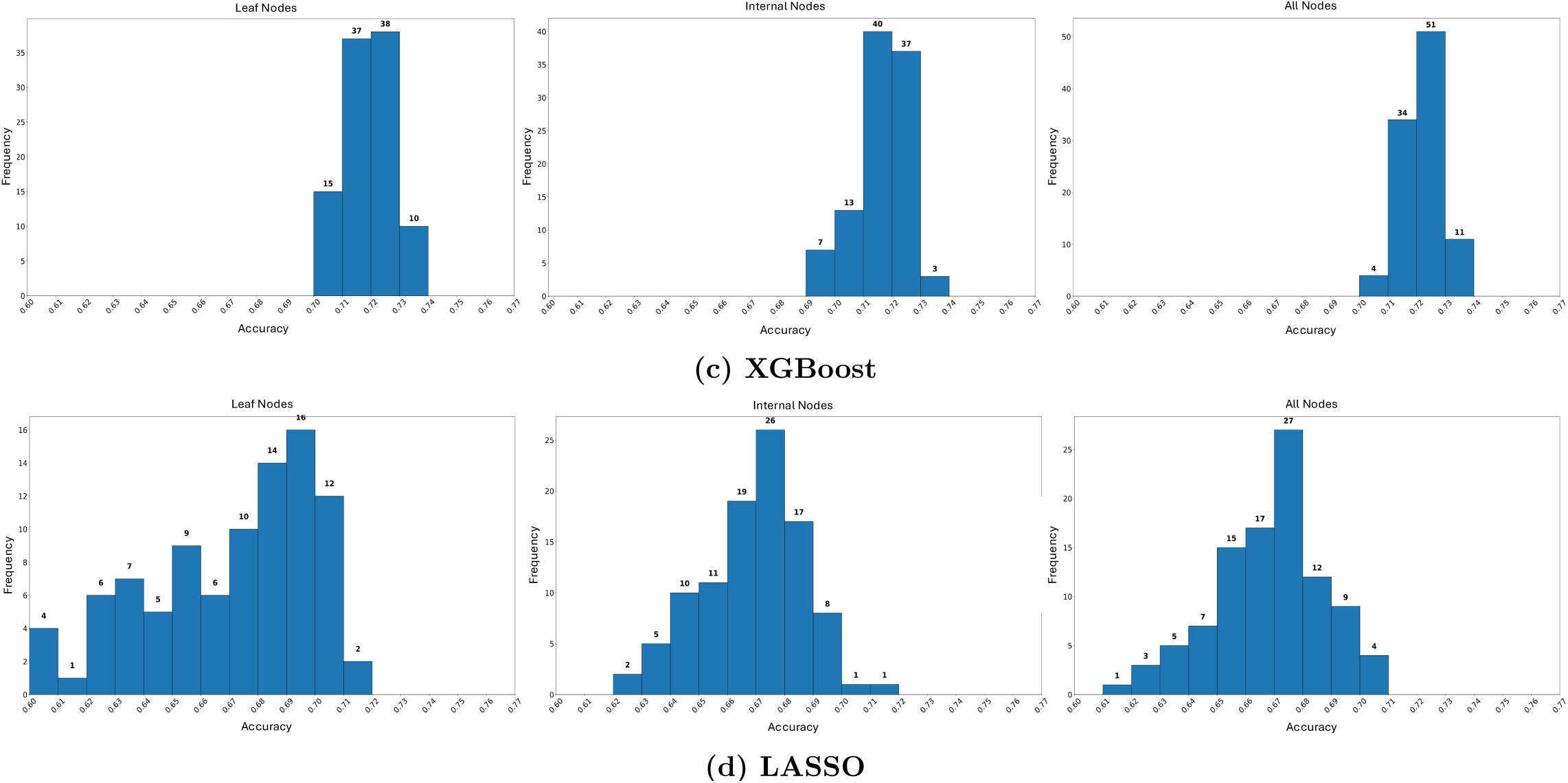
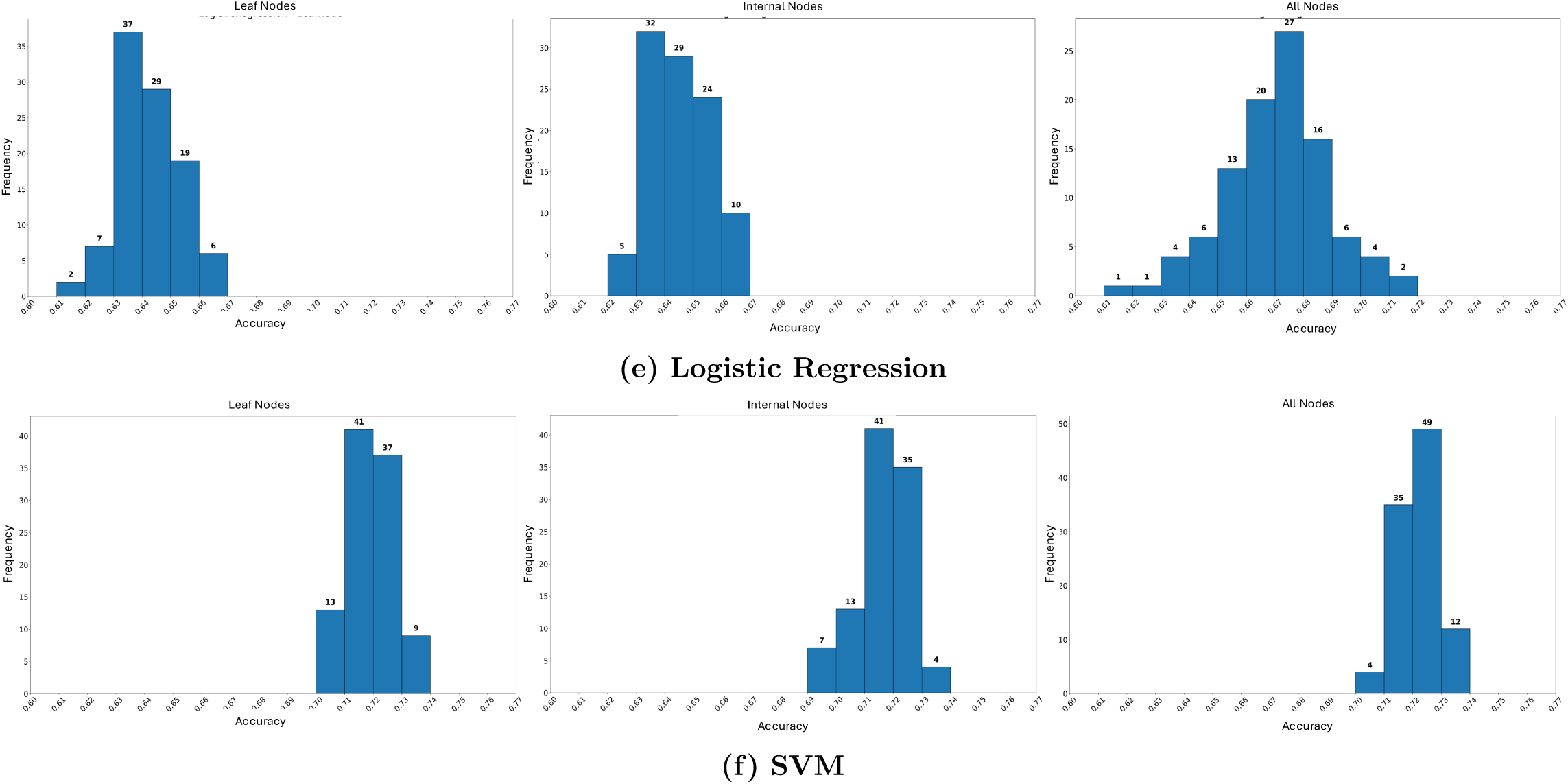
Comparative analysis of classification accuracy with CART as a base model. Panels show: (a) RF, (b) GBM, (c) XGBoost, (d) LASSO, (e) Logistic Regression, and (f) SVM for leaf nodes, internal nodes, and all nodes.

**Fig. 8:**
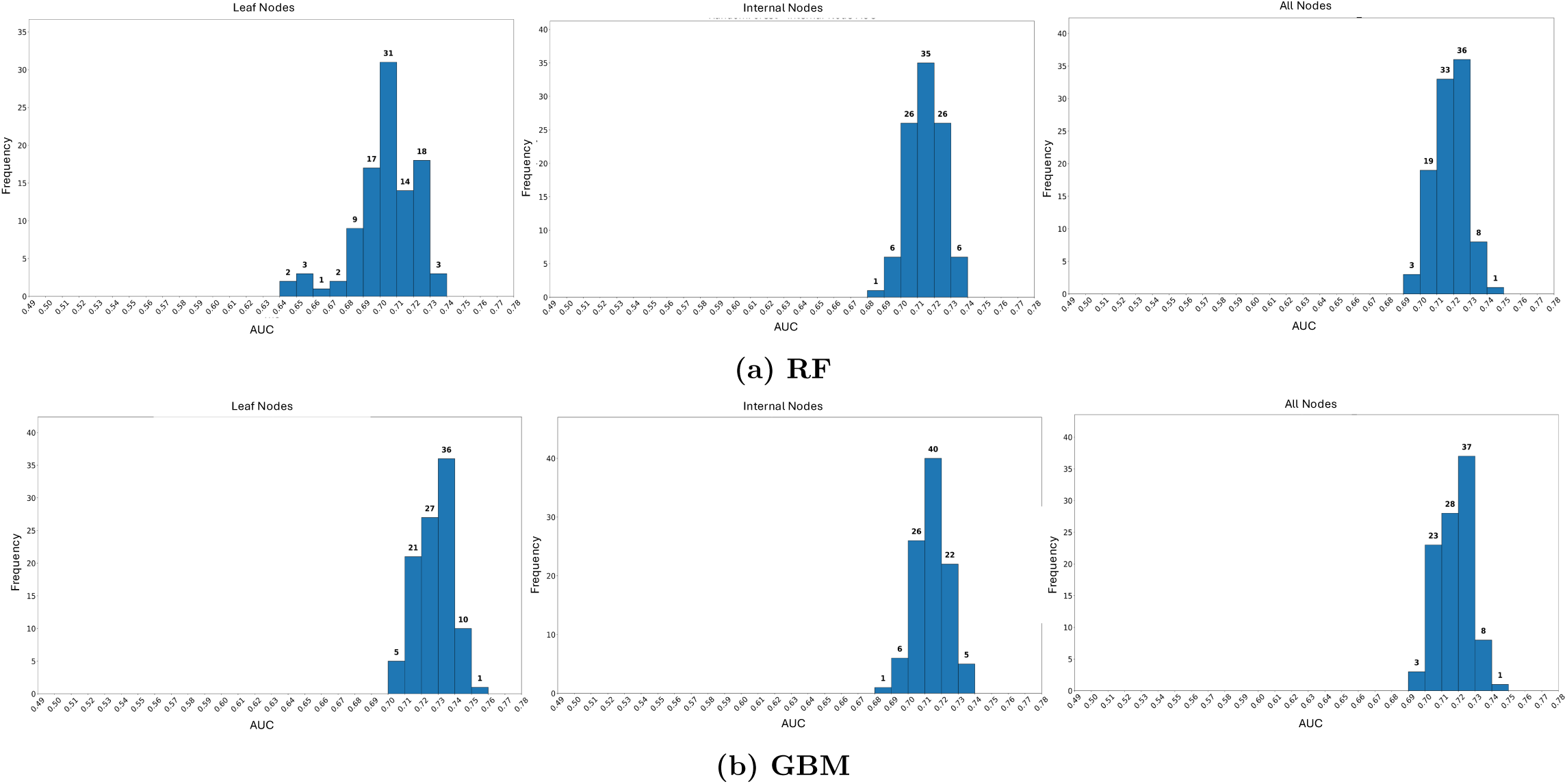
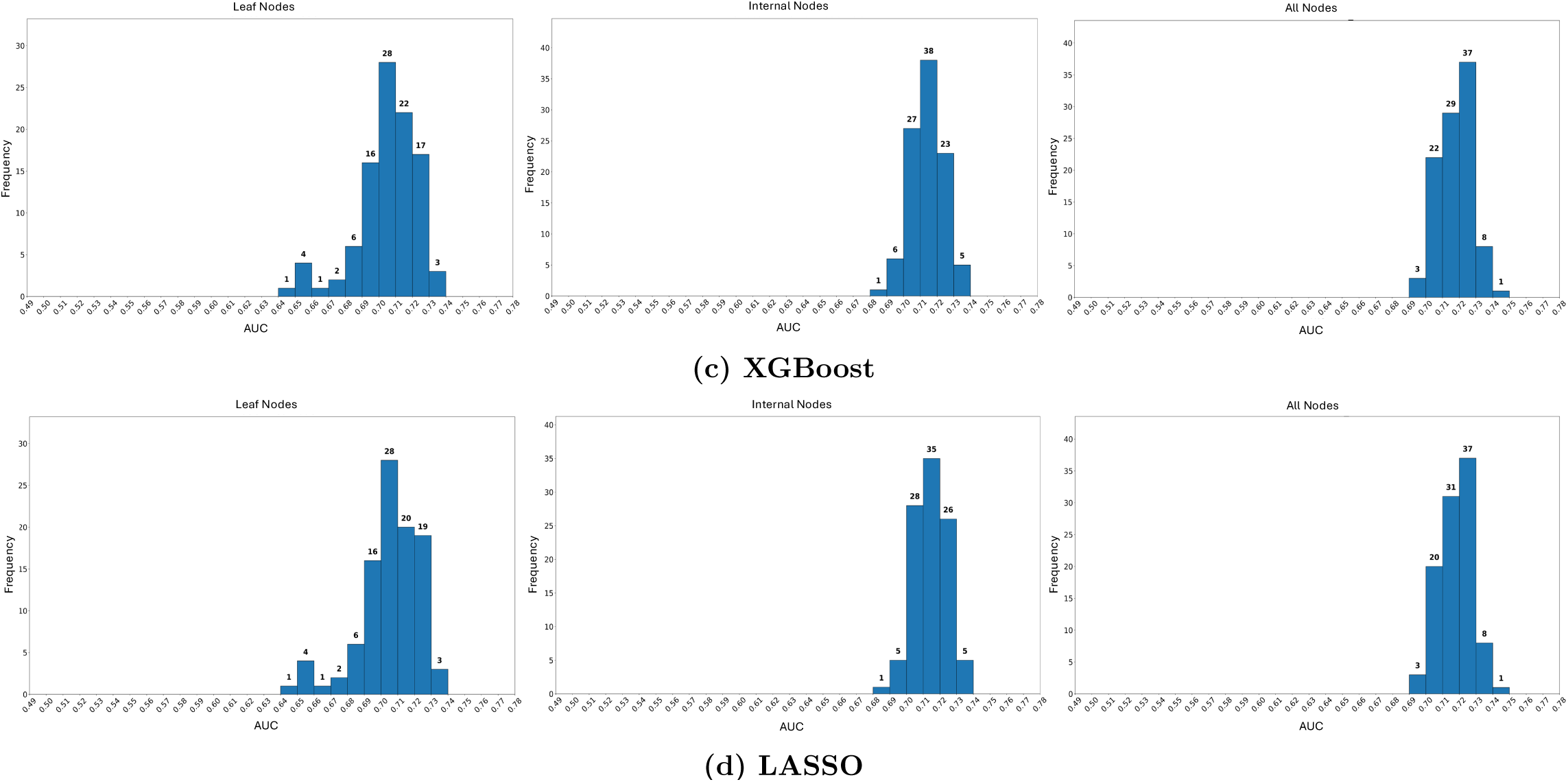
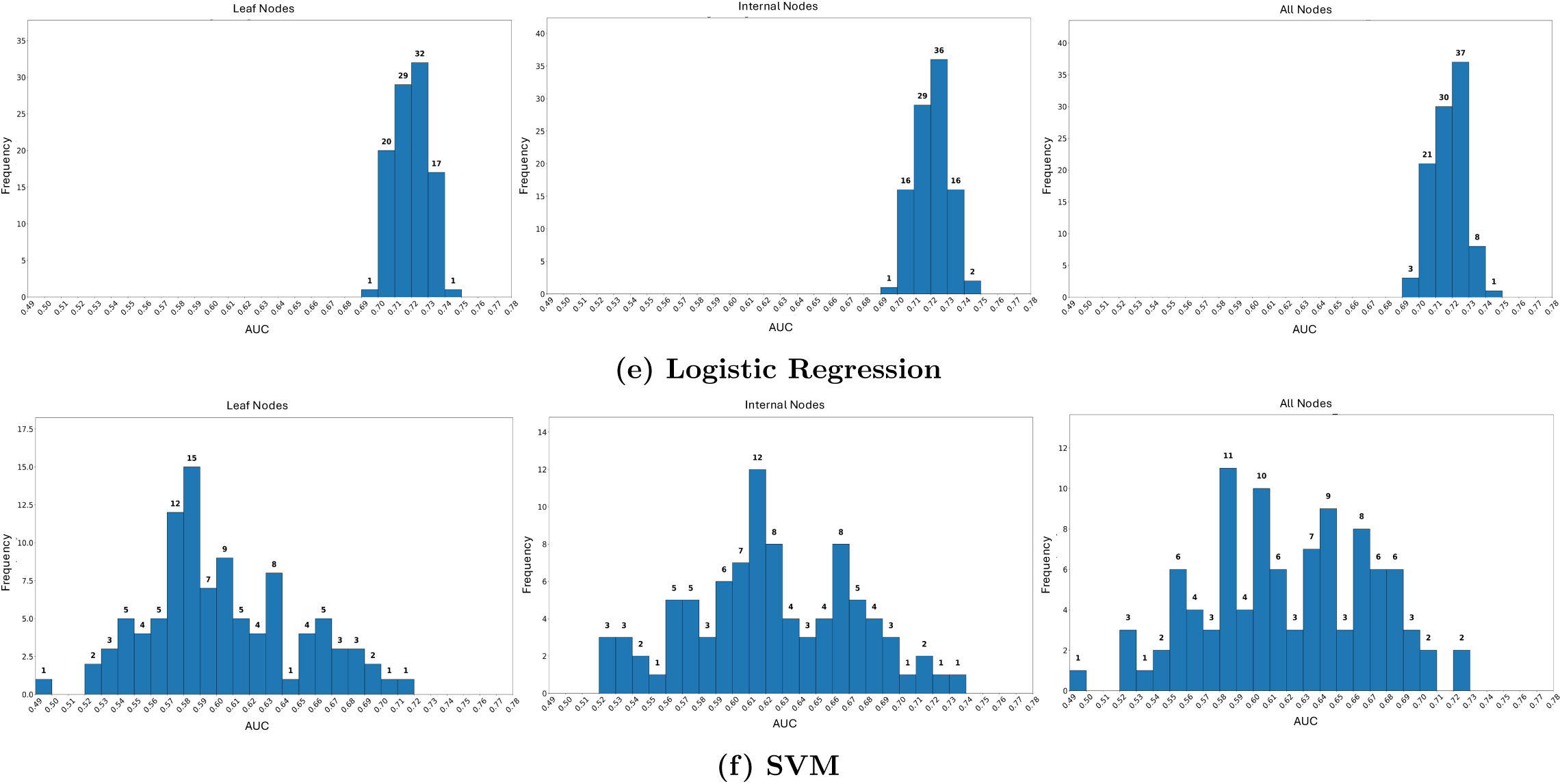
Comparative analysis of AUC with CART as a base model. Panels show: (a) RF, (b) GBM, (c) XGBoost, (d) LASSO, (e) Logistic Regression, and (f) SVM for leaf nodes, internal nodes, and all nodes.

For feature selection, this approach achieved a competitive central tendency, often comparable to Grid Search, but with moderately increased variability in Figures 6. For the SVM (Figures 6(f)), however, Feature Selection produced the most concentrated AUC distribution, suggesting that reducing the feature space is particularly vital for margin-based classifiers to mitigate the noise inherent in high-dimensional GWAS data.

Finally, for CART-based dummy variables (Complete Tree), this novel approach exhibited the greatest dispersion and the most pronounced signs of multimodality across all models. The increased variability is likely attributable to structural instability in the decision trees used to generate the dummy variables. While some configurations achieved peak performance, the overall distributions suggest less consistent results than direct optimization of the raw feature space.

#### 4.3.2 Comparative Classification Summary

The quantitative summary of discriminative performance (Tables 1 to 3) underscores that Grid Search Optimization outperformed other approaches for maximizing class separability in this dataset.

**Table 2:** Overview of Classification Accuracy using Grid Search Approach.

| Algorithm | Mean (%) | Medium (%) | Maximum (%) | Minimum (%) |
| --- | --- | --- | --- | --- |
| GBoost | 72.89 | 72.91 | 75.38 | 70.15 |
| LASSO | 64.37 | 64.31 | 66.54 | 62.02 |
| Logistic Regression | 64.34 | 64.18 | 66.54 | 62.06 |
| Random Forest | 72.19 | 72.19 | 73.89 | 70.16 |
| XGBoost | 72.41 | 72.40 | 73.94 | 70.78 |
| SVM | 71.99 | 72.06 | 73.59 | 70.37 |
| CART | 71.50 | 71.53 | 73.31 | 69.73 |

**Table 3:** Overview of Classification Accuracy using Feature Selection Approach.

| Algorithm | Mean (%) | Medium (%) | Maximum (%) | Minimum (%) |
| --- | --- | --- | --- | --- |
| GBoost | 72.36 | 72.41 | 74.52 | 70.03 |
| LASSO | 64.18 | 64.09 | 66.48 | 61.88 |
| Logistic Regression | 64.26 | 64.19 | 66.60 | 61.94 |
| Random Forest | 72.07 | 72.10 | 73.77 | 69.18 |
| XGBoost | 72.28 | 72.31 | 74.28 | 70.27 |
| SVM | 71.45 | 71.57 | 73.69 | 68.12 |
| CART | 71.88 | 71.94 | 73.20 | 69.40 |

The classification accuracy results in (Tables 1 to 3) compare the performance of the Brute-Force, Grid Search, and Feature Selection approaches across seven classifiers. In these three approaches, CART-derived dummy variables are not used; instead, the models are evaluated using the original feature representation under different model-selection strategies. Overall, the ensemble-based classifiers achieved the strongest performance, while the linear models produced comparatively lower accuracy. In particular, GBoost, XGBoost, Random Forest, SVM, and CART generally achieved mean accuracies in the range of approximately 70-73%, whereas LASSO and Logistic Regression remained near 64%. This pattern suggests that nonlinear and ensemble-based classifiers are better suited for capturing the predictive structure in the original feature space than linear models.

Among the three approaches, the Grid Search approach produced the best overall classification accuracy. GBoost with Grid Search achieved the highest mean accuracy of 72.89% and the highest maximum accuracy of 75.38%. XGBoost and Random Forest also performed strongly under Grid Search, with mean accuracies of 72.41% and 72.19%, respectively. These results indicate that hyperparameter tuning improves model performance, particularly for ensemble-based classifiers. The improvement of SVM from 64.97% under the Brute-Force approach to 71.99% under Grid Search further shows that SVM is highly sensitive to parameter selection.

The Feature Selection approach also achieved competitive performance. GBoost achieved a mean accuracy of 72.36%, XGBoost achieved 72.28%, and Random Forest achieved 72.07%. Although these values are slightly lower than the corresponding Grid Search results, they remain close to the best-performing models. This indicates that Feature Selection is able to remove redundant or less informative variables while preserving most of the predictive signal. Therefore, Feature Selection provides a useful trade-off between classification performance and model simplicity.

In contrast, LASSO and Logistic Regression consistently produced the lowest mean accuracies across all three approaches. Their mean accuracies remained around 64%, suggesting that linear decision boundaries are insufficient for fully capturing the predictive structure of the original feature set. CART performed better than the linear models but was generally weaker than the ensemble methods, which is expected because a single decision tree is more sensitive to data variation than ensemble models such as GBoost, XGBoost, and Random Forest.

Overall, the results demonstrate that ensemble-based classifiers are the most effective for this classification task when dummy variables are not used. GBoost with Grid Search provides the strongest overall performance, while XGBoost and Random Forest show stable and competitive results across all three approaches. The results also suggest that Feature Selection can retain much of the predictive information while reducing model complexity, making it a practical alternative when interpretability and reduced dimensionality are important.

#### 4.3.3 Comparative AUC Summary

The AUC results in (Tables 4 to 6) provide a threshold-independent evaluation of the Brute-Force, Grid Search, and Feature Selection approaches. Unlike accuracy, which depends on a fixed decision threshold, AUC measures how well each classifier separates the two classes across all possible thresholds. Therefore, the AUC results are useful for determining whether a model has strong discriminative ability, even when its classification accuracy is similar to that of other models.

**Table 4:**
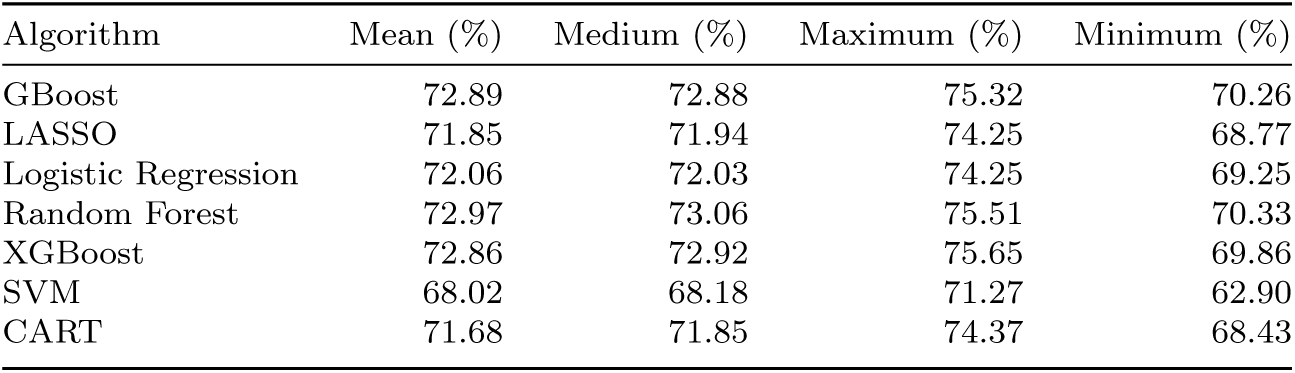
Overview of AUC using the Brute-Force Approach.

| Algorithm | Mean (%) | Medium (%) | Maximum (%) | Minimum (%) |
| --- | --- | --- | --- | --- |
| GBoost | 72.89 | 72.88 | 75.32 | 70.26 |
| LASSO | 71.85 | 71.94 | 74.25 | 68.77 |
| Logistic Regression | 72.06 | 72.03 | 74.25 | 69.25 |
| Random Forest | 72.97 | 73.06 | 75.51 | 70.33 |
| XGBoost | 72.86 | 72.92 | 75.65 | 69.86 |
| SVM | 68.02 | 68.18 | 71.27 | 62.90 |
| CART | 71.68 | 71.85 | 74.37 | 68.43 |

**Table 5:**
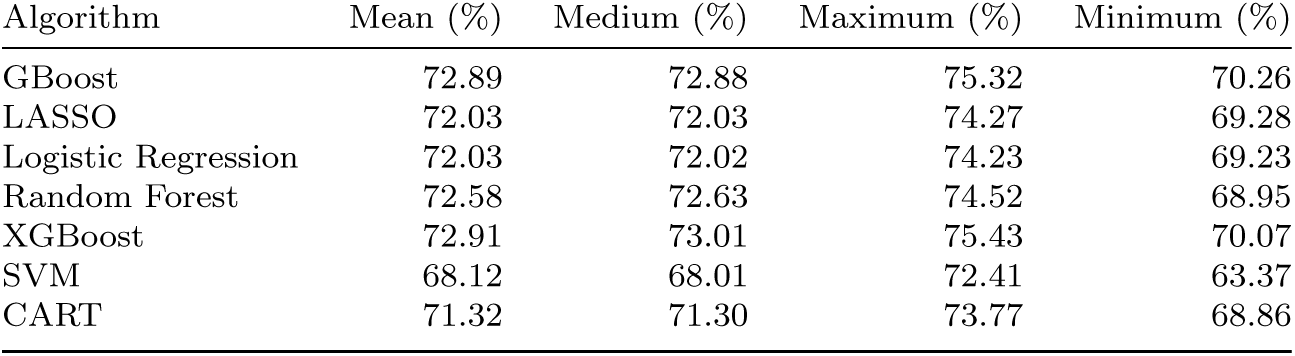
Overview of AUC using the Grid Search Approach.

| Algorithm | Mean (%) | Medium (%) | Maximum (%) | Minimum (%) |
| --- | --- | --- | --- | --- |
| GBoost | 72.89 | 72.88 | 75.32 | 70.26 |
| LASSO | 72.03 | 72.03 | 74.27 | 69.28 |
| Logistic Regression | 72.03 | 72.02 | 74.23 | 69.23 |
| Random Forest | 72.58 | 72.63 | 74.52 | 68.95 |
| XGBoost | 72.91 | 73.01 | 75.43 | 70.07 |
| SVM | 68.12 | 68.01 | 72.41 | 63.37 |
| CART | 71.32 | 71.30 | 73.77 | 68.86 |

**Table 6:** Overview of AUC using Feature Selection Approach.

| Algorithm | Mean (%) | Medium (%) | Maximum (%) | Minimum (%) |
| --- | --- | --- | --- | --- |
| GBoost | 72.04 | 72.12 | 74.60 | 68.82 |
| LASSO | 71.59 | 71.65 | 74.15 | 69.05 |
| Logistic Regression | 71.23 | 71.27 | 74.41 | 67.93 |
| Random Forest | 72.20 | 72.39 | 74.77 | 69.53 |
| XGBoost | 71.94 | 72.03 | 74.40 | 69.35 |
| SVM | 66.17 | 67.16 | 71.36 | 50.91 |
| CART | 71.58 | 71.61 | 73.76 | 68.92 |

Overall, the AUC results show that the ensemble-based classifiers provide stronger discrimination than the linear models. GBoost, XGBoost, and Random Forest generally achieve higher AUC values, indicating that these models are better able to rank case and control samples. This pattern is consistent with the accuracy results, where ensemble-based methods also produced the strongest classification performance. The stronger AUC values suggest that their performance advantage is not limited to fixed-threshold classification, but also reflects improved class separation.

The Grid Search approach shows the strongest overall AUC performance among the three approaches. This indicates that hyperparameter tuning improves not only accuracy but also the ranking ability of the classifiers. In particular, models such as GBoost and XGBoost benefit from parameter optimization because their performance depends strongly on tuning parameters such as learning rate, tree depth, number of estimators, and regularization. Therefore, the Grid Search results suggest that careful model tuning is important for achieving better discrimination between the two classes.

The Feature Selection approach also produces competitive AUC results. Although its AUC values may be slightly lower than those obtained using Grid Search, the results remain close to the best-performing models. This indicates that Feature Selection successfully removes less informative or redundant variables while retaining the most important predictive information. As a result, Feature Selection provides a useful balance between model performance and reduced feature complexity.

In contrast, LASSO and Logistic Regression show weaker AUC performance compared with the ensemble-based classifiers. This suggests that linear models have limited ability to capture the nonlinear relationships and interaction effects present in the data. Even when their accuracy values appear reasonable, lower AUC values indicate weaker ranking ability and poorer separation between the two classes.

Overall, the AUC results support the conclusion drawn from the accuracy analysis: ensemble-based classifiers, particularly GBoost and XGBoost, are the most effective models for this classification task. The Grid Search approach provides the strongest overall performance, while Feature Selection offers a competitive and more compact alternative. The combined accuracy and AUC results indicate that the best-performing models improve both hard-label classification and threshold-independent class discrimination.

#### 4.3.4 Overall Interpretation

Overall, the accuracy and AUC results show that ensemble-based classifiers provide the strongest performance across the Brute-Force, Grid Search, and Feature Selection approaches. GBoost, XGBoost, and Random Forest generally achieve higher accuracy and stronger AUC values, indicating that these models improve both fixed-threshold classification and threshold-independent class separation. Grid Search gives the best overall performance, showing the benefit of hyperparameter tuning, while Feature Selection remains competitive by reducing less informative variables without greatly reducing performance. In contrast, LASSO and Logistic Regression perform weaker, suggesting that linear models are less effective for capturing the predictive structure of the data.

Although the baseline approaches provide useful predictive performance, they do not explicitly expose the decision structure learned by CART. The use of dummy variables is introduced to make this structure more interpretable and analytically useful. By converting CART decision paths or nodes into dummy-variable features, each selected rule can be represented as a separate binary indicator. This allows the downstream classifiers to use tree-derived subgroup information while also enabling us to identify which CART-based rules contribute most to classification performance. Therefore, the dummy-variable approach shifts the analysis from only measuring prediction accuracy to understanding how specific decision rules and subgroup patterns influence the final classification outcome.

### 4.4 Methodological Efficiency and Computational Latency

To extend the baseline analysis, we further introduce CART-based dummy-variable approaches. While the previous Brute-Force, Grid Search, and Feature Selection results evaluate classifier performance using the original feature representation, these additional approaches are designed to examine whether the structural information learned by CART can improve interpretability and provide more detailed insight into the classification process. Specifically, the decision tree is first converted into a set of dummy variables, where each dummy variable represents a node-based decision rule. This allows the analysis to move beyond overall predictive performance and investigate how different parts of the CART structure contribute to classification.

To provide a comprehensive evaluation, two additional node-based approaches are considered:

#### 4.4.1 Evaluation using the Leaf Nodes

In this approach, only the leaf nodes are considered for evaluation. Initially, CART-based dummy variables are generated from the complete decision tree. For the analysis, only the dummy variables corresponding to the leaf nodes are retained, as shown in Figure 4, where the leaf nodes are highlighted in red. In this figure, the dummy leaf nodes are indexed, and the evaluation is performed using the proposed dummy-variable-based approach.

#### 4.4.2 Evaluation using the Internal Nodes

In the second approach, the analysis is restricted to internal nodes. A complete set of CART-based dummy variables is first generated from the full decision tree. Subsequently, only the dummy variables corresponding to internal nodes are retained for analytical evaluation, as illustrated in Figure 4, where the internal nodes are highlighted in blue. In this figure, the dummy variables associated with internal nodes are systematically indexed, and the evaluation is conducted using the proposed methodology.

#### 4.4.3 Analysis of the All Nodes, Leaf Nodes, and Internal Nodes

We evaluate classification performance across three CART-derived feature representations: leaf nodes, internal nodes, and all nodes. The accuracy results are summarized in (Tables 7, to 9), while the corresponding AUC results are presented in (Tables 10 to 12). Together, these tables provide a comparative view of how different portions of the CART structure contribute to downstream classification performance.

**Table 7:** Overview of Classification Accuracy using Leaf Nodes with CART as Base Model.

| Algorithm | Mean (%) | Medium (%) | Maximum (%) | Minimum (%) |
| --- | --- | --- | --- | --- |
| GBoost | 72.60 | 72.58 | 74.26 | 70.63 |
| LASSO | 66.20 | 67.23 | 71.70 | 52.02 |
| Logistic Regression | 64.30 | 64.22 | 66.70 | 61.62 |
| Random Forest | 71.06 | 71.04 | 73.37 | 69.56 |
| XGBoost | 71.94 | 71.94 | 73.73 | 70.14 |
| SVM | 71.94 | 71.91 | 73.73 | 70.14 |

**Table 8:** Overview of Classification Accuracy using Internal Nodes with CART as Base Model.

| Algorithm | Mean (%) | Medium (%) | Maximum (%) | Minimum (%) |
| --- | --- | --- | --- | --- |
| GBoost | 72.74 | 72.69 | 74.32 | 71.07 |
| LASSO | 64.29 | 64.13 | 66.49 | 62.07 |
| Logistic Regression | 64.48 | 64.30 | 66.54 | 62.39 |
| Random Forest | 72.33 | 72.36 | 73.73 | 70.98 |
| XGBoost | 72.74 | 72.69 | 74.32 | 71.07 |
| SVM | 72.36 | 72.39 | 74.10 | 70.96 |

**Table 9:** Overview of Classification Accuracy using All Nodes with CART as Base Model.

| Algorithm | Mean (%) | Medium (%) | Maximum (%) | Minimum (%) |
| --- | --- | --- | --- | --- |
| GBoost | 72.17 | 72.13 | 73.62 | 70.63 |
| LASSO | 66.87 | 67.04 | 70.91 | 61.95 |
| Logistic Regression | 67.10 | 67.23 | 71.31 | 61.95 |
| Random Forest | 72.18 | 72.15 | 73.62 | 69.95 |
| XGBoost | 72.17 | 72.13 | 73.62 | 70.63 |
| SVM | 72.16 | 72.11 | 73.62 | 70.63 |

**Table 10:**
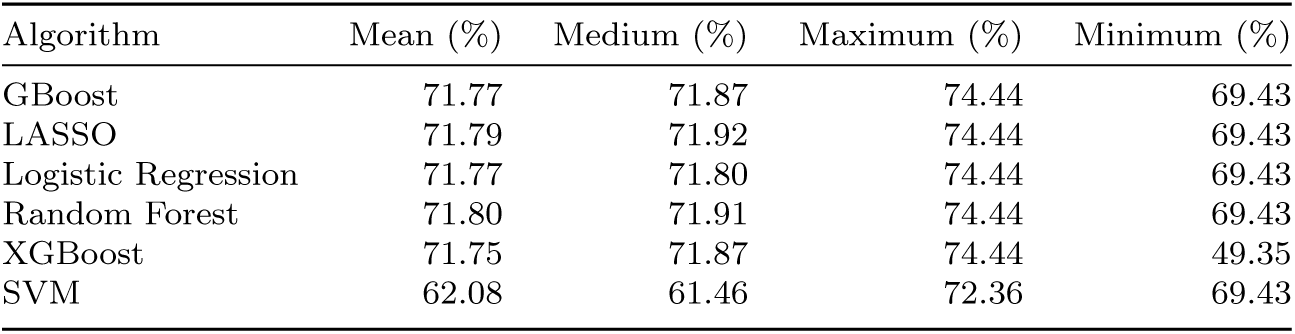
Overview of Classification AUC using All Nodes with CART as Base Model.

**Table 11:** Overview of Classification AUC using Leaf Nodes with CART as Base Model.

| Algorithm | Mean (%) | Medium (%) | Maximum (%) | Minimum (%) |
| --- | --- | --- | --- | --- |
| GBoost | 72.79 | 72.84 | 75.01 | 70.24 |
| LASSO | 70.52 | 70.69 | 73.80 | 64.01 |
| Logistic Regression | 71.97 | 71.99 | 74.23 | 69.22 |
| Random Forest | 70.38 | 70.59 | 73.67 | 64.45 |
| XGBoost | 70.51 | 70.68 | 73.82 | 64.78 |
| SVM | 60.39 | 59.32 | 71.74 | 49.53 |

**Table 12:** Overview of Classification AUC using Internal Nodes with CART as Base Model.

| Algorithm | Mean (%) | Medium (%) | Maximum (%) | Minimum (%) |
| --- | --- | --- | --- | --- |
| GBoost | 71.44 | 71.52 | 73.88 | 68.57 |
| LASSO | 71.47 | 71.54 | 73.87 | 68.57 |
| Logistic Regression | 72.06 | 72.03 | 74.25 | 69.25 |
| Random Forest | 71.49 | 71.57 | 73.90 | 68.57 |
| XGBoost | 71.44 | 71.50 | 73.88 | 68.57 |
| SVM | 59.91 | 61.56 | 73.24 | 30.49 |

The leaf-node representation in Tables 7 provides strong performance for several classifiers, with GBoost achieving a mean accuracy of 72.60% and XGBoost and SVM both achieving mean accuracies of 71.94%. In terms of AUC, Table 11 shows that the leaf-node approach produces the highest AUC for GBoost, with a mean AUC of 72.79% and a maximum AUC of 75.01%. This suggests that leaf nodes capture useful terminal decision patterns that can improve class discrimination, especially for boosting-based models.

The internal-node representation in (Table 8) achieves slightly stronger accuracy for several ensemble-based classifiers. GBoost and XGBoost both obtain mean accuracies of 72.74%, while Random Forest and SVM achieve mean accuracies of 72.33% and 72.36%, respectively. These results indicate that internal nodes preserve important intermediate decision-rule information from the CART structure. However, the AUC results in (Table 12) show that the internal-node representation does not uniformly dominate the leaf-node representation. Logistic Regression achieves the highest internal-node mean AUC of 72.06%, while GBoost and XGBoost obtain mean AUC values of 71.44%. SVM shows a lower mean AUC of 59.91%, indicating weaker ranking stability for this representation.

The all-node representation combines both leaf-node and internal-node information. As shown in (Table 9), this representation produces stable accuracy across the ensemble-based classifiers, with GBoost, Random Forest, XGBoost, and SVM all achieving mean accuracies around 72%. This indicates that using all CART-derived nodes provides a balanced feature representation. However, the all-node approach does not always produce the highest accuracy compared with the internal-node representation. For example, GBoost and XGBoost achieve higher mean accuracy using internal nodes than using all nodes.

In terms of AUC, (Table 10) shows that the all-node representation provides highly consistent AUC values for most classifiers, with GBoost, LASSO, Logistic Regression, Random Forest, and XGBoost all obtaining mean AUC values around 72%. This consistency suggests that the all-node representation provides stable threshold-independent discrimination across models.

Overall, the results indicate that no single CART-derived representation dominates across both accuracy and AUC for every classifier. Internal nodes provide the strongest accuracy for several ensemble models, while leaf nodes produce the strongest AUC for GBoost. The all-node representation offers the most balanced and stable performance across classifiers by integrating both terminal and intermediate decision-rule information. Therefore, the all-node representation can be considered the most robust overall feature representation, while leaf-node and internal-node representations provide useful complementary perspectives on how different parts of the CART structure contribute to classification performance.

### 4.5 Computational Time Analysis

We further evaluate the computational cost associated with different modeling strategies, including Brute-Force evaluation, Grid Search optimization, Feature Selection, and CART-based dummy variable construction, as shown in (Tables 13 and 14). The total execution time is measured in terms of real (wall-clock), user (CPU processing), and system time across all experimental runs.

**Table 13:** Computation time comparison between All-Nodes (CART-Dummy Tree), Leaf-Nodes, and Internal-Nodes.

| Algorithm | Leaf-Nodes Run Time<br>(minutes) | Internal-Nodes Run Time<br>(minutes) | All-Nodes Run Time<br>(minutes) |
| --- | --- | --- | --- |
| GBoost | 138.81 | 86.75 | 1954.01 |
| LASSO | 140.83 | 86.89 | 1962.16 |
| Logistic Regression | 149.89 | 109.62 | 1960.49 |
| Random Forest | 150.52 | 110.02 | 1955.56 |
| XGBoost | 309.27 | 103.51 | 2382.50 |
| SVM | 164.01 | 168.45 | 1974.05 |

**Table 14:**
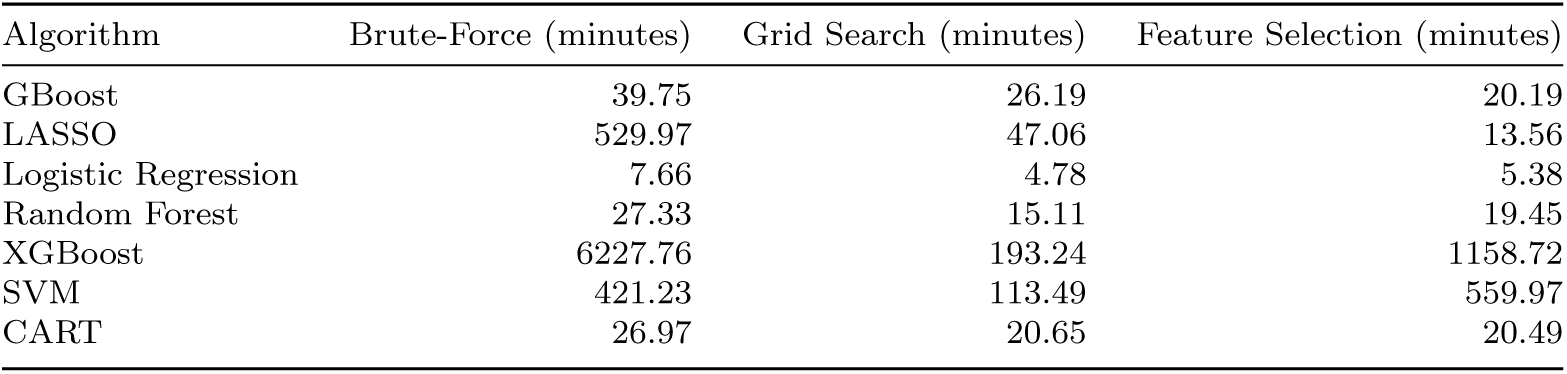
Computation time comparison between Brute-Force, Grid Search, and Feature Selection.

| Algorithm | Brute-Force (minutes) | Grid Search (minutes) | Feature Selection (minutes) |
| --- | --- | --- | --- |
| GBoost | 39.75 | 26.19 | 20.19 |
| LASSO | 529.97 | 47.06 | 13.56 |
| Logistic Regression | 7.66 | 4.78 | 5.38 |
| Random Forest | 27.33 | 15.11 | 19.45 |
| XGBoost | 6227.76 | 193.24 | 1158.72 |
| SVM | 421.23 | 113.49 | 559.97 |
| CART | 26.97 | 20.65 | 20.49 |

Among the evaluated approaches, grid search exhibits the highest computational overhead due to exhaustive hyperparameter exploration. This significantly increases runtime, particularly for ensemble models such as Random Forest, Gradient Boosting, and XGBoost, where multiple trees and parameter combinations must be evaluated.

Feature Selection introduces additional preprocessing cost; however, it reduces the dimensionality of the input space, leading to more efficient downstream training. As a result, feature selection achieves a balance between computational efficiency and predictive performance.

The Brute-Force approach, while computationally expensive due to repeated evaluations across multiple training-validation splits, benefits from simpler model configurations and avoids the overhead of hyperparameter tuning.

The proposed CART-based dummy variable approach introduces an initial over-head for tree construction and feature generation. However, once the dummy features are created, subsequent model training becomes more efficient, particularly for linear models. Furthermore, this approach improves model stability and reduces variability across repeated runs, offsetting its initial computational cost.

Overall, ensemble methods incur higher computational costs compared to linear models but consistently provide superior predictive performance. These results highlight a trade-off between computational efficiency and model accuracy, where optimization strategies and feature engineering play a critical role in balancing performance and runtime.

Although the CART-based feature engineering approach introduces additional pre-processing time, it significantly reduces model variance and improves stability across repeated runs. This trade-off is particularly beneficial in high-dimensional genomic datasets, where capturing feature interactions is critical for predictive performance.

## 5 Discussion

The results demonstrate that model-selection strategy and feature representation both influence classification performance. In the baseline analysis, where CART-derived dummy variables were not used, the Brute-Force, Grid Search, and Feature Selection approaches showed that ensemble-based classifiers generally outperformed linear models. GBoost, XGBoost, and Random Forest consistently achieved stronger accuracy and AUC values, indicating that nonlinear and interaction-based learning methods are better suited for capturing the predictive structure of the data. Among the baseline approaches, Grid Search provided the strongest overall performance, showing that hyperparameter optimization improves both fixed-threshold accuracy and threshold-independent class discrimination. However, this improvement comes with increased computational cost, particularly when repeated across many training splits and classifiers.

The CART-based dummy-variable framework provides an additional layer of analysis by converting the learned decision-tree structure into interpretable node-based features. This representation allows downstream classifiers to use decision-rule information derived from CART while also making it possible to compare the contribution of leaf nodes, internal nodes, and all nodes. The leaf-node results show that terminal decision regions contain useful class-separation information, with strong AUC performance for GBoost. The internal-node results show slightly stronger accuracy for several ensemble classifiers, suggesting that intermediate decision rules preserve important subgroup structure before the final terminal classification. The all-node representation combines both terminal and intermediate decision information and provides the most balanced and stable performance across classifiers.

Across the CART-derived representations, no single node type dominates every classifier and metric. Internal nodes provide strong accuracy for several ensemble models, while leaf nodes provide strong AUC performance in selected cases. The all-node representation does not always produce the highest value for every classifier, but it offers the most consistent performance by integrating both local terminal patterns and broader internal decision rules. This makes the all-node approach a robust compromise between predictive performance and interpretability.

Overall, the findings suggest that advanced optimization methods such as Grid Search can improve predictive performance, but they also increase computational burden. In contrast, the CART-based dummy-variable framework offers a practical and interpretable feature-engineering strategy. It preserves complex decision-tree interactions, enables efficient downstream classification, and provides insight into how different parts of the tree contribute to prediction. Therefore, combining CART-derived dummy variables with ensemble-based classifiers provides a useful balance between accuracy, AUC, stability, and interpretability.

## 6 Conclusion

This study evaluated multiple classification strategies using both baseline feature representations and CART-derived dummy-variable representations. The baseline results showed that ensemble-based classifiers, especially GBoost, XGBoost, and Random Forest, achieved stronger accuracy and AUC performance than linear models such as LASSO and Logistic Regression. Grid Search produced the best overall baseline performance, confirming the value of hyperparameter tuning, although at a higher computational cost.

The CART-derived dummy-variable analysis further demonstrated that tree-based decision rules can be transformed into useful predictive features. Leaf-node, internal-node, and all-node representations each captured different aspects of the CART structure. Leaf nodes reflected terminal decision patterns, internal nodes captured intermediate subgroup rules, and all nodes combined both sources of information. Among these, the all-node representation provided the most stable and balanced performance across classifiers, while leaf-node and internal-node representations offered complementary advantages for specific models and metrics.

In conclusion, the results indicate that CART-based dummy-variable feature engineering is a practical approach for improving interpretability while maintaining competitive classification performance. Ensemble-based classifiers remain the most effective downstream models, and the all-node CART representation provides a robust feature space for balancing predictive accuracy, AUC-based discrimination, and model interpretability.

## 7 Discussion

## Ethics approval and consent to participate

All participants in this study provided written informed consent, and the local institutional review boards approved the study.

## Consent for publication

Not Applicable

## Availability of data and materials

The datasets used and analyzed during the current study are available from the corresponding author on reasonable request.

## Authors’ contributions

Jinyoung Byun (JB), Dheeman Saha (DS), Younghun Han (YH), and Christopher I. Amos (CIA) conceived the work. YH prepared the genetic data. DS conducted the data analyses. JB, DS, and CIA interpreted the results. JB and DS drafted the manuscript, and all authors revised and contributed to the intellectual content of the article. All authors approved the final version of the article, including the authorship list.

## Acknowledgements

Not Applicable

## Appendix A Supplementary Tables

**Table A1:**
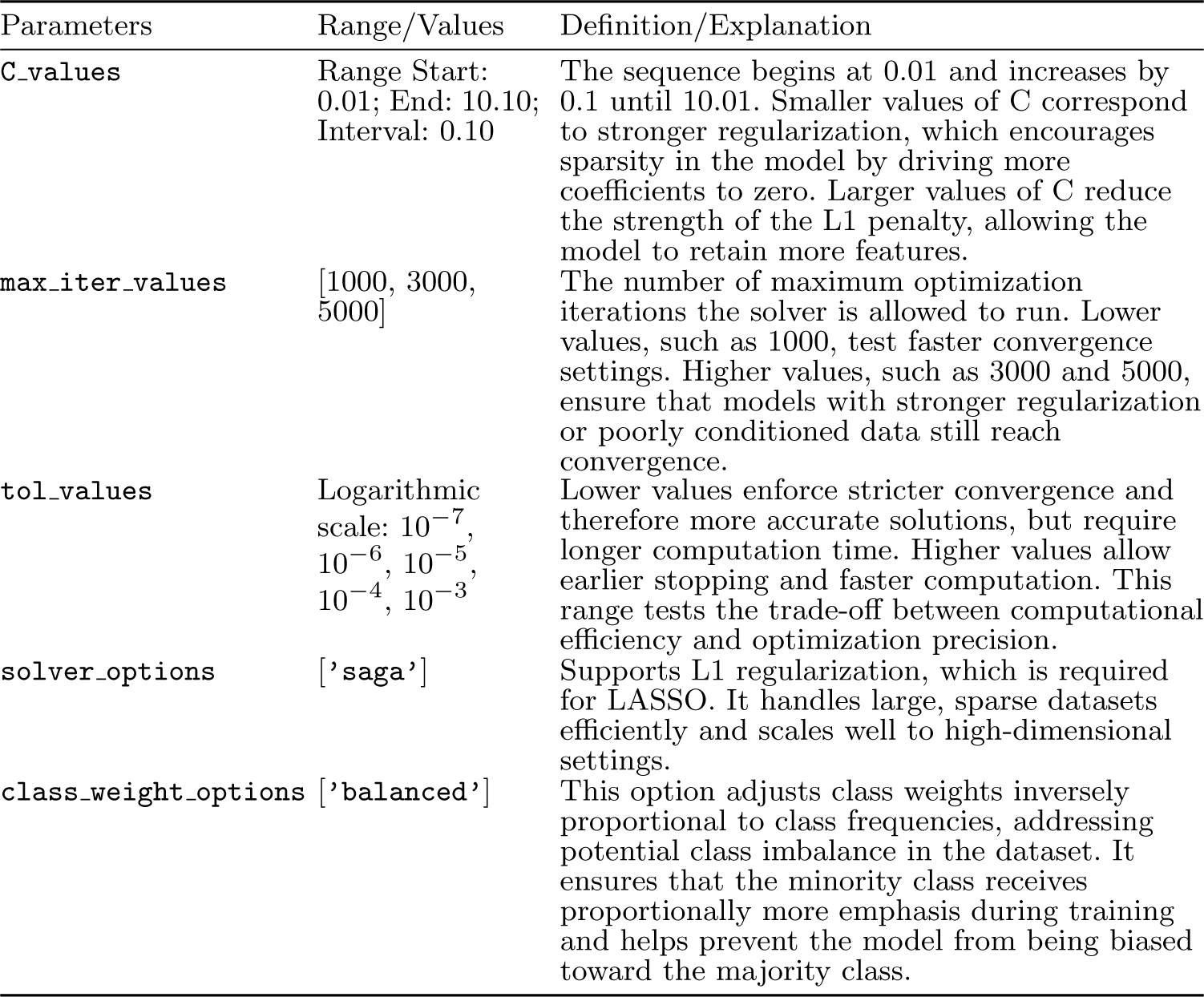
Parameters associated with the Least Absolute Shrinkage and Selection Operator (LASSO) modeling.

| Parameters | Range/Values | Definition/Explanation |
| --- | --- | --- |
| **C_values** | Range Start: 0.01; End: 10.10; Interval: 0.10 | The sequence begins at 0.01 and increases by 0.1 until 10.01. Smaller values of C correspond to stronger regularization, which encourages sparsity in the model by driving more coefficients to zero. Larger values of C reduce the strength of the L1 penalty, allowing the model to retain more features. |
| **max_iter_values** | [1000, 3000, 5000] | The number of maximum optimization iterations the solver is allowed to run. Lower values, such as 1000, test faster convergence settings. Higher values, such as 3000 and 5000, ensure that models with stronger regularization or poorly conditioned data still reach convergence. |
| **tol_values** | Logarithmic scale: $10^{-7}$, $10^{-6}$, $10^{-5}$, $10^{-4}$, $10^{-3}$ | Lower values enforce stricter convergence and therefore more accurate solutions, but require longer computation time. Higher values allow earlier stopping and faster computation. This range tests the trade-off between computational efficiency and optimization precision. |
| **solver_options** | ['saga'] | Supports L1 regularization, which is required for LASSO. It handles large, sparse datasets efficiently and scales well to high-dimensional settings. |
| **class_weight_options** | ['balanced'] | This option adjusts class weights inversely proportional to class frequencies, addressing potential class imbalance in the dataset. It ensures that the minority class receives proportionally more emphasis during training and helps prevent the model from being biased toward the majority class. |

**Table A2:**
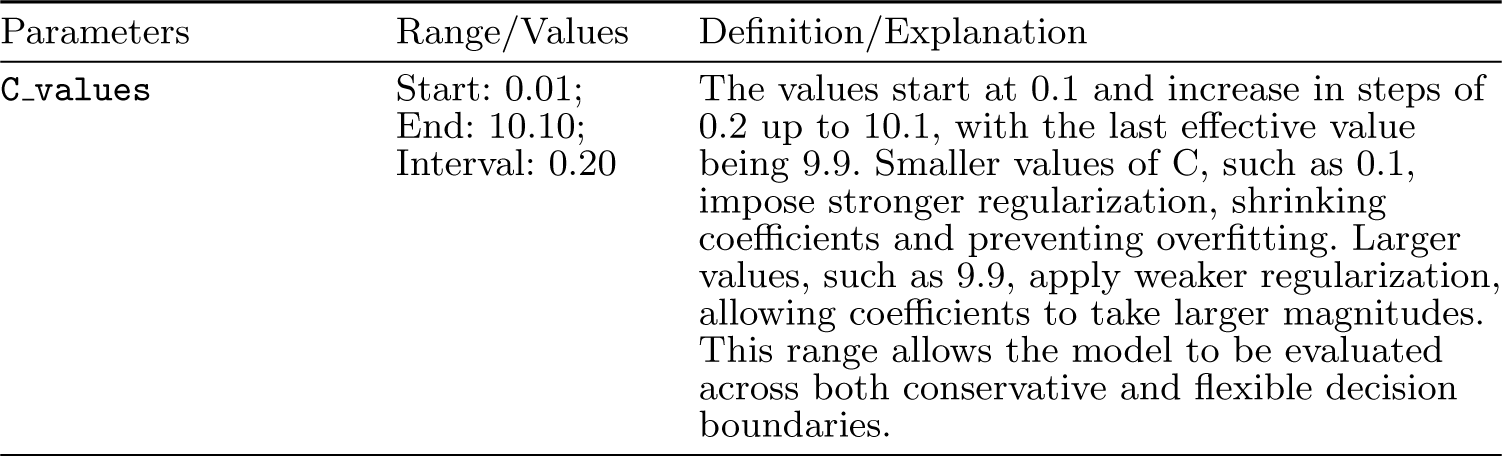
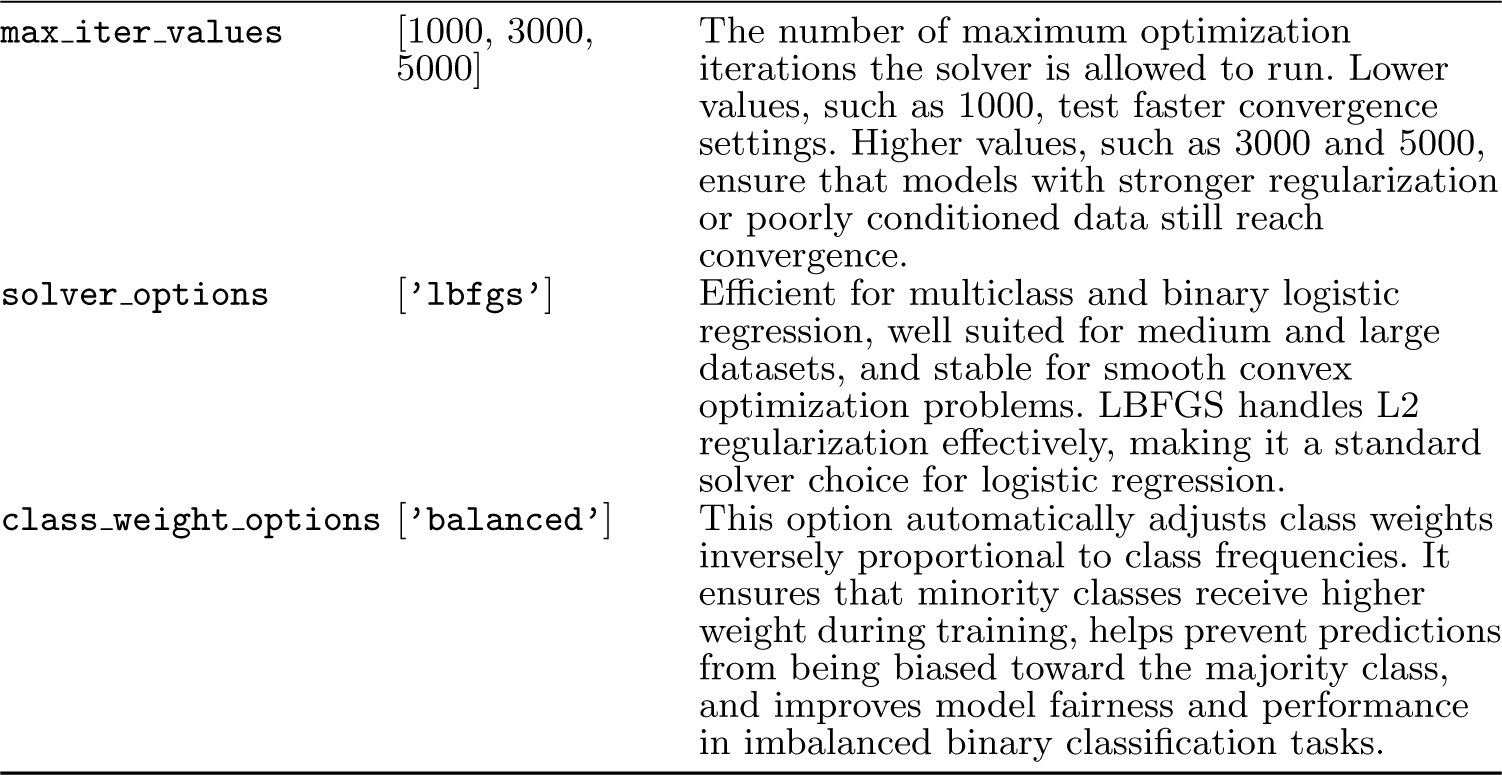
Parameters associated with Logistic Regression modeling.

| Parameters | Range/Values | Definition/Explanation |
| --- | --- | --- |
| **C_values** | Start: 0.01; End: 10.10; Interval: 0.20 | The values start at 0.1 and increase in steps of 0.2 up to 10.1, with the last effective value being 9.9. Smaller values of C, such as 0.1, impose stronger regularization, shrinking coefficients and preventing overfitting. Larger values, such as 9.9, apply weaker regularization, allowing coefficients to take larger magnitudes. This range allows the model to be evaluated across both conservative and flexible decision boundaries. |

**Table A2:** Parameters associated with Logistic Regression modeling. Continued.
| Parameters | Range/Values | Definition/Explanation |
| --- | --- | --- |
| `max_iter_values` | [1000, 3000, 5000] | The number of maximum optimization iterations the solver is allowed to run. Lower values, such as 1000, test faster convergence settings. Higher values, such as 3000 and 5000, ensure that models with stronger regularization or poorly conditioned data still reach convergence. |
| `solver_options` | ['lbfgs'] | Efficient for multiclass and binary logistic regression, well suited for medium and large datasets, and stable for smooth convex optimization problems. LBFGS handles L2 regularization effectively, making it a standard solver choice for logistic regression. |
| `class_weight_options` | ['balanced'] | This option automatically adjusts class weights inversely proportional to class frequencies. It ensures that minority classes receive higher weight during training, helps prevent predictions from being biased toward the majority class, and improves model fairness and performance in imbalanced binary classification tasks. |

**Table A3:** Parameters associated with Support Vector Machine (SVM) modeling.

| Parameters | Range/Values | Definition/Explanation |
| --- | --- | --- |
| `C_values` | [0.1, 1, 10] | This is the regularization parameter controlling the trade-off between maximizing the margin and minimizing classification errors. A value of 0.1 applies strong regularization, producing a wider margin and greater tolerance for misclassification, but with potential underfitting risk. A value of 1 provides a standard balance between margin width and classification fidelity. A value of 10 applies weaker regularization, producing a tighter fit to the data and increasing the risk of overfitting. |
| `kernels` | ['linear', 'rbf', 'poly'] | These represent three commonly used SVM kernel functions. The linear kernel is appropriate for linearly separable or high-dimensional sparse data. The RBF kernel is widely used for capturing nonlinear relationships by mapping data to higher-dimensional space. The polynomial kernel captures structured nonlinear interactions and is useful for datasets with polynomial-like boundaries. |
| `gammas` | ['scale', 0.1] | This parameter controls the kernel coefficient for RBF and polynomial kernels. The 'scale' option is the recommended automatic setting and adapts to the dataset. The fixed value 0.1 introduces additional flexibility and allows the model to fit more complex boundaries. This combination evaluates both adaptive and explicit gamma settings. |

**Table A3:** Parameters associated with Support Vector Machine (SVM) modeling. Continued.
| Parameters | Range/Values | Definition/Explanation |
| --- | --- | --- |
| **degrees** | [3] | This parameter specifies the degree of the polynomial kernel. A lower degree of 3 is the standard and most widely used choice, providing a balanced nonlinear representation. |
| **class_weights** | [None, 'balanced'] | This parameter determines how the SVM handles class imbalance. None gives all classes equal weight. The 'balanced' option assigns weights inversely proportional to class frequencies, giving more importance to the minority class and helping reduce majority-class bias. |

**Table A4:** Parameters associated with Classification and Regression Trees (CART) modeling.

| Parameters | Range/Values | Definition/Explanation |
| --- | --- | --- |
| **max_depth_values** | Start: 1; End: 21; Interval: 1 | This range specifies the maximum depth allowed for each decision tree. Smaller depths, such as 1–5, produce shallow, highly constrained trees, reducing overfitting and yielding simpler, more interpretable decision boundaries. Larger depths, such as 10–20, allow the tree to grow more complex, capturing deeper feature interactions at the risk of overfitting. This comprehensive range allows evaluation across minimal to highly expressive tree structures. |
| **min_samples_split_values** | Start: 2; End: 11; Interval: 1 | This defines the minimum number of samples required to split an internal node. A value of 2 leads to the most flexible trees, allowing many splits. Higher values, such as 8–10, enforce stricter conditions for splitting, producing more conservative trees. This parameter helps control tree complexity and prevent overfitting, especially in noisy datasets. |
| **min_samples_leaf_values** | Start: 1; End: 6; Interval: 1 | This specifies the minimum number of samples that must be present in a leaf node. A leaf size of 1 provides the most granular splits but may lead to overfitting. Larger leaf sizes, such as 4–5, produce smoother, more generalized decision regions. Tuning this parameter balances model flexibility and stability. |
| **criterion_options** | ['gini', 'entropy'] | These are the available impurity metrics used to evaluate split quality. Gini impurity is computationally faster and is the default choice in CART implementations. Entropy, also known as information gain, is based on information theory and is often preferred when interpretability in terms of information gain is desired. |

**Table A5:** Parameters associated with Random Forest modeling.

| Parameters | Range/Values | Definition/Explanation |
| --- | --- | --- |
| `n_estimators` | [100, 300, 500] | The number of trees in the forest. Higher values provide more stable results but require longer training time. |
| `max_depths` | [4, 6] | The maximum depth of each decision tree. Smaller depths produce simpler trees and reduce overfitting. |
| `min_samples_splits` | [2, 5] | Minimum number of samples required to split an internal node. A value of 2 allows flexible trees, while increasing to 5 reduces overfitting by requiring more data before splitting. |
| `min_samples_leafs` | [1, 3] | Minimum number of samples required at a leaf node. A value of 1 allows pure leaves, while 3 produces smoother decision boundaries and reduces overfitting. |
| `min_impurity_decrease` | [0.0] | A node is split only if the split reduces impurity by at least this threshold. A value of 0.0 indicates no restriction. |
| `ccp_alphas` | [0.0] | Cost-complexity pruning parameter. A value of 0.0 indicates no pruning, while higher values prune more aggressively. |
| `min_weight_fraction_leaf` | [0.0] | Minimum weighted fraction of samples required at a leaf node. A value of 0.0 disables this restriction. |
| `criterion_list` | [gini, entropy] | Split-quality measure. Gini represents Gini impurity, while entropy is the alternative impurity criterion. |
| `max_features_list` | [sqrt] | Number of features used to search for the best split. The sqrt option uses the square root of the number of features. |
| `max_leaf_nodes_list` | [None] | Limits the number of leaf nodes. None allows trees to grow until stopping parameters are reached. |
| `bootstrap_options` | [True] | Indicates whether bootstrap sampling is used. True means each tree is trained on a random sample with replacement. |
| `oob_score` | [False] | Out-of-bag scoring. If True, validation accuracy is estimated without requiring a separate validation set. |
| `n_jobs` | [None] | Controls parallel processing. None uses a single CPU core, while -1 uses all available cores. |
| `random_state` | [42] | Seed for reproducibility, ensuring consistent results across runs. |
| `verbose` | [0] | Controls logging output during training. A value of 0 indicates silent mode. |
| `warm_starts` | [False] | If True, allows forests to grow incrementally. False trains a fresh forest each time. |
| `class_weight` | [None] | Class-weighting strategy. None gives equal weight to all classes. |
| `max_samples_list` | [None] | Used only when bootstrap=True. None draws a full bootstrap sample. |

**Table A6:** Parameters associated with XGBoost modeling.

| Parameters | Range/Values | Definition/Explanation |
| --- | --- | --- |
| **n_estimators** | [100, 300, 500] | Number of boosted trees. Lower values provide faster training, while higher values increase model capacity. |
| **learning_rates** | [0.05, 0.1] | Controls how much each new tree contributes to the ensemble. Smaller values produce conservative updates, while larger values increase learning speed. |
| **max_depths** | [3, 5] | Maximum depth of each tree. Smaller depths reduce overfitting, while larger depths allow more complex feature interactions. |
| **subsamples** | [0.8, 1.0] | Fraction of samples used for each boosting iteration. A value of 0.8 introduces stochasticity, while 1.0 uses all samples. |

**Table A7:**
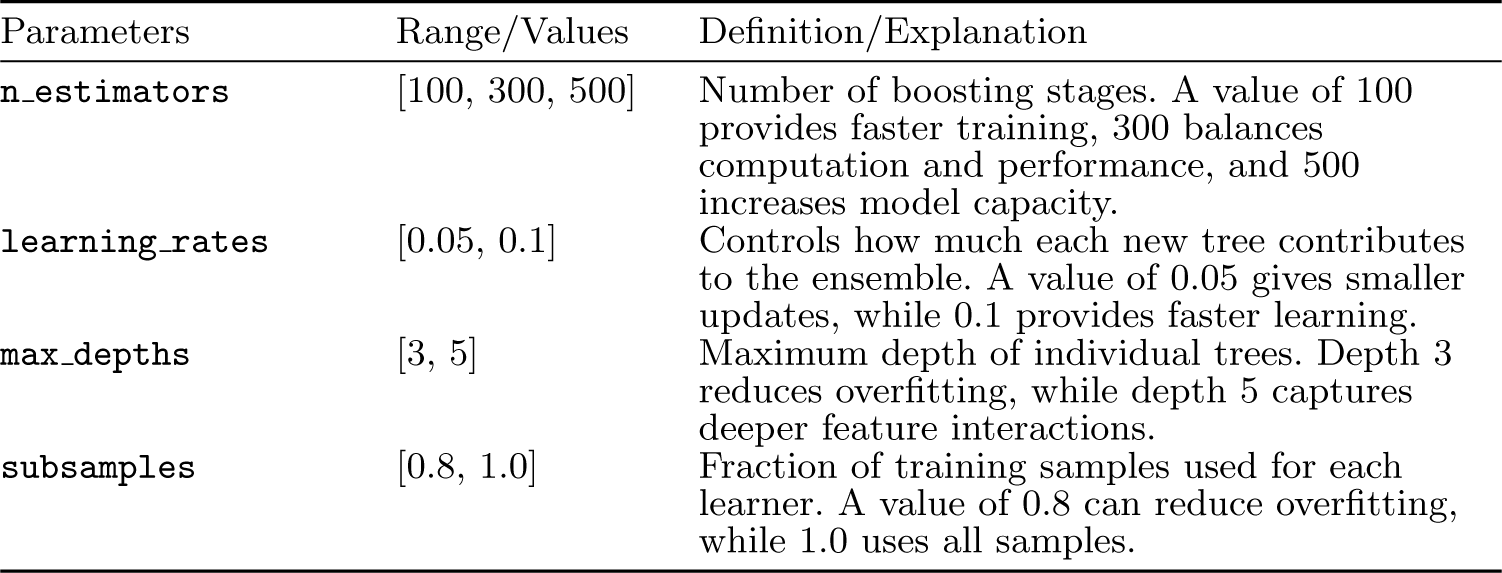
Parameters associated with Gradient Boosting modeling.

